# Chronic nanoplastic exposure causes size-dependent cytotoxicity and ANGPTL4 induction in human kidney cells

**DOI:** 10.64898/2026.10.08.757574

**Authors:** Hayden L. Gillings, Darling M. Rojas-Canales, Soon Wei Wong, Timothy Burton, Jonathan M. Gleadle, Melanie MacGregor

**Affiliations:** Flinders Institute for Nanoscale Science and Technology, College of Science and Engineering, Flinders University, Sturt Road, Bedford Park, South Australia, 5042, Australia; Flinders Health and Medical Research Institute, College of Medicine and Public Health, Flinders University, Sturt Road, Bedford Park, South Australia, 5042, Australia; Nano and Microplastics Research Consortium, Flinders University, Sturt Road, Bedford Park, South Australia, 5042, Australia; Department of Renal Medicine, Flinders Medical Centre, Flinders Drive, Bedford Park, South Australia, 5042, Australia

**Keywords:** nanoplastics, kidney, proximal tubule, polystyrene, long-term exposure, *ANGPTL4*

## Abstract

Nanoplastics (NPs) are widespread environmental contaminants, yet most toxicity studies rely on short-term, single-dose exposures that poorly reflect daily human exposure. The kidney, particularly the proximal tubule, is a likely site of repeated NP exposure given its role in filtration and reabsorption, but chronic exposure effects are largely unexplored. Here, a long-term *in vitro* model was developed in which human proximal tubule (HK-2) cells were continuously exposed to 20 nm or 100 nm carboxylated polystyrene NPs (100 µg/mL) over 12 days. Continuous 20 nm NP exposure caused progressive loss of viability whereas 100 nm NPs left viable cell numbers largely unaffected. RNA-sequencing showed 20 nm NPs drove extensive, time-dependent transcriptional dysregulation (1,548 differentially expressed genes at day 6, rising to 4,431 at day 9), far exceeding the minimal alterations in gene expression (35 genes) caused by 100 nm NPs. Cell-cycle analysis and pathway enrichment revealed KRAS pathway activation and S-phase arrest in 20 nm-treated cells, with *ANGPTL4* the most upregulated gene. Validation with qRT-PCR confirmed increased *ANGPTL4*, *HMOX1* and *FN1* expression, genes implicated in renal fibrosis and chronic kidney disease. These findings show that continuous, size-dependent NP exposure elicits progressive cytotoxic and transcriptional responses in kidney proximal tubule cells and indicate ANGPTL4 as a candidate early biomarker of NP-induced renal stress.

## 1. Introduction

Exposure to microplastics (MPs) and nanoplastics (NPs) is becoming an ubiquitous part of daily life, with numerous studies identifying accumulation, predominantly MPs, within the human body.^1, 2^ MPs and NPs have been observed in virtually all tissues, organs and organ systems, and bodily fluids,^1^ including a recent study from Massardo et al.^3^ which identified 66 MPs of 1 µM and larger in 10 samples of healthy kidney tissue and urine.

Although substantial research on the health effects of NPs exposure is underway, the vast majority of studies are only investigating short-term exposure, typically 24-48 hours, with a single dosage of NPs. Recent reports estimate the average human is exposed to approximately 113,000-126,000 MPs per year,^4^ or potentially as low as 52 individual MPs or all the way up to 168,000 MPs per day.^5^ These estimates, and the broad range they span, highlight the difficulties not only with determining current exposure levels, but also studying the impact of exposure with environmentally relevant concentrations. Further still, the investigation of the risks of NPs exposure within the kidney, and the renal system as a whole, is limited.^6^

Given the function of the renal system in filtering the blood and tubular reabsorption, it is likely that the kidney experiences frequent exposure to NPs. Filtration within the kidney is predominantly managed by the nephron, which consists of the glomerulus and renal tubules.^7, 8^ The passive filtration ability of the glomerulus is governed by the glomerular filtration barrier (GFB), which has an established limit of approximately 6-8 nm, though in kidney disease disruption of the GFB can result in filtration of particles above the established size limit.^9, 10^ When functioning normally, the GFB utilises endocytic pathways to filter through larger particles for proximal tubule uptake.^11, 12^ The filtrate is reabsorbed in the renal tubules, where the proximal tubule plays a dominant role in maintaining the body’s homeostasis, by recovering glucose, proteins, and electrolytes that have been filtered from blood by the glomerulus.^7, 8, 13^ Of particular concern is that this exposure to NPs might contribute to the development or progression of chronic kidney disease (CKD), a condition affecting an estimated 10% of the global population and one of the leading causes of morbidity and mortality worldwide.^14, 15^ Renal interstitial fibrosis is recognised as the common final pathway underlying CKD progression irrespective of the initiating insult, with the extent of fibrosis directly correlating with loss of renal function.^15, 16^ Given the proximal tubule’s sustained, direct contact with filtered NPs, and the established role of tubular epithelial cell stress and dysfunction in initiating fibrotic remodelling, it is critical to establish whether chronic NP exposure represents an unrecognised contributor to fibrosis-driven CKD.

Current kidney-focused NPs studies are limited in assessing potential cytotoxicity and acute response with short-term exposure times, often only for 24 hours,^17–19^ though sometimes spanning as short as 12 hours,^20^ and as long as 48 hours.^21, 22^ Across these studies, proximal tubule human kidney-2 (HK-2) cells, human renal tubular epithelial cells (HKC), and human embryonic kidney (293T) cells were exposed to PS NPs between 20 nm and 500 nm, at concentrations between 1 µg/mL and 800 µg/mL, and size-and concentration-dependent effects were broadly identified.^17–22^ The greatest cytotoxicity was typically identified in the smallest 20 nm NPs,^17, 20^ and the severity of the cytotoxicity was correlated with increasing concentrations.^18, 19, 21^ The minimum concentration of NPs required to induce significant cytotoxicity varies between studies, with one study identifying toxicity within 24 hours with as little as 5 µg/mL of NPs,^17^ while another study only saw toxicity after exposure to concentrations of 400 µg/mL and higher.^18^ Numerous studies have also identified increased rates of oxidative stress,^21^ increased generation of reactive oxygen species (ROS),^17^ and reduced mitochondrial function.^18, 20^

In a previous study, we assessed the acute impacts of polystyrene (PS), polyethylene (PE) and poly(methyl methacrylate) (PMMA) NPs on HK-2 cells within 24-hours of exposure.^23^ The HK-2 cells were exposed to concentrations of 0.2-200 µg/mL, differing particle sizes (15-100 nm) and different suspension formulations, in order to study dose-, size- and additive-dependent effects.^23^ For the PS NPs, significant cytotoxicity was limited to the highest 200 µg/mL concentration, though no change to viability was seen in the 20 nm and 100 nm PS NPs supplied without additives.^23^ Interestingly, dysregulation of the G0/G1 and S cell cycle phases was observed in the smallest size groups (15 nm and 20 nm).^23^

To help better understand the impact of systemic exposure to NPs and examine for the potential toxicity of more chronic exposure, we developed an *in vitro* assay to model continuous, repeated NPs exposure over a 12-day period. HK-2 cells were exposed to 100 µg/mL doses of 20 nm (7.80×10^12^ particles/mL) and 100 nm (1.36×10^11^ particles/mL) carboxyl (-COOH) surface modified PS NPs from the time of seeding to maximise sustained NP-cell interactions. This model was used to compare the effects of 20 nm and 100 nm PS NPs on cellular morphology, cell cycle progression, and transcriptional responses.

The work shows that continuous, size-dependent NP exposure elicits progressive cytotoxic and transcriptional responses in kidney proximal tubule cells and indicate ANGPTL4 as a candidate early biomarker of NP-induced renal stress.

## 2. Methods

### 2.1 Nanoplastics dilution and treatment

Carboxylated (COOH-) PS NPs of 20 nm and 100 nm in diameter were purchased from Thermo Fisher Scientific. All NP stocks, dilutions and treatments were briefly vortexed and sonicated for 15 minutes in a water bath sonicator (Soniclean Digital Benchtop Ultrasonic Cleaner 80TD) immediately prior to dilution or addition to cells to facilitate homogenisation of the suspensions. Both NP suspensions were diluted in 1x PBS (Gibco) to a concentration of 1 mg/mL. Immediately prior to cell treatment, both NP dilutions were further diluted 1:10 into cell culture media for a final concentration of 100 µg/mL. Details of the NPs and the concentration used for treatment are provided in Table S1, where the number of particles/mL and particles/cell were calculated as follows:

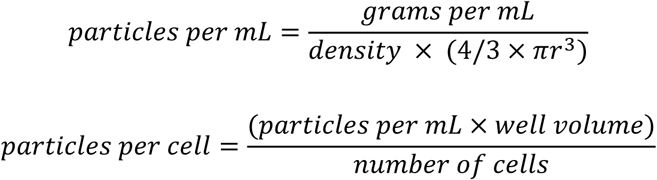

In these equations, *r* (nm) refers to the mean radius of the NPs.

### 2.2 Cell culture and nanoplastics exposure

A long-term NPs exposure study was performed following a modified version of the method outlined by Saadati, et al..^24^ HK-2 cells (ATCC CRL-2190) were cultured in complete media consisting of 1:1 Dulbecco’s Modified Eagle Medium (DMEM) and Ham’s F-12 (Gibco), 10 mM HEPES (Gibco), 10% fetal bovine serum (FCS; Bovogen Biologicals) and 1X penicillin, streptomycin and glutamine mix (Gibco) at 37°C with 5% CO_2_. Cells were seeded in T-75 flasks at 1.2 × 10^6^ cells/flask in 12 mL of complete media containing either the vehicle control (10% 1X PBS) or the 20 nm or 100 nm PS NPs diluted to a final concentration of 100 µg/mL. The flasks were monitored every 24-hours and then split after 72-hours of continuous NPs exposure. Cell numbers and viability were assessed via the Countess 3 FL Automated Cell Counter (Invitrogen) with a 1:1 dilution of Trypan blue (Gibco), and 2 replicates analysed per sample. New T-75 flasks were seeded with 1.2×10^6^ cells/flask with the respective control or treatment media, aliquots of the cell suspension were then collected for the cell cycle (1.0 × 10^6^ cells), western blot (1.0 × 10^6^ cells), and RNA-sequencing analyses (0.1 × 10^6^ cells). Any remaining cell suspension was spun down at 300 x g for 5 minutes, and the supernatant removed before immediate storage at -80°C. A graphical summary of the experiment is provided in Fig. 1a. In separate experiments, HK-2 cells were treated with 100 µg/mL and 1 mg/mL concentrations of the 20 nm and 100 nm PS NPs for 24-hours prior to collection for western blot analysis.

**Fig. 1.**
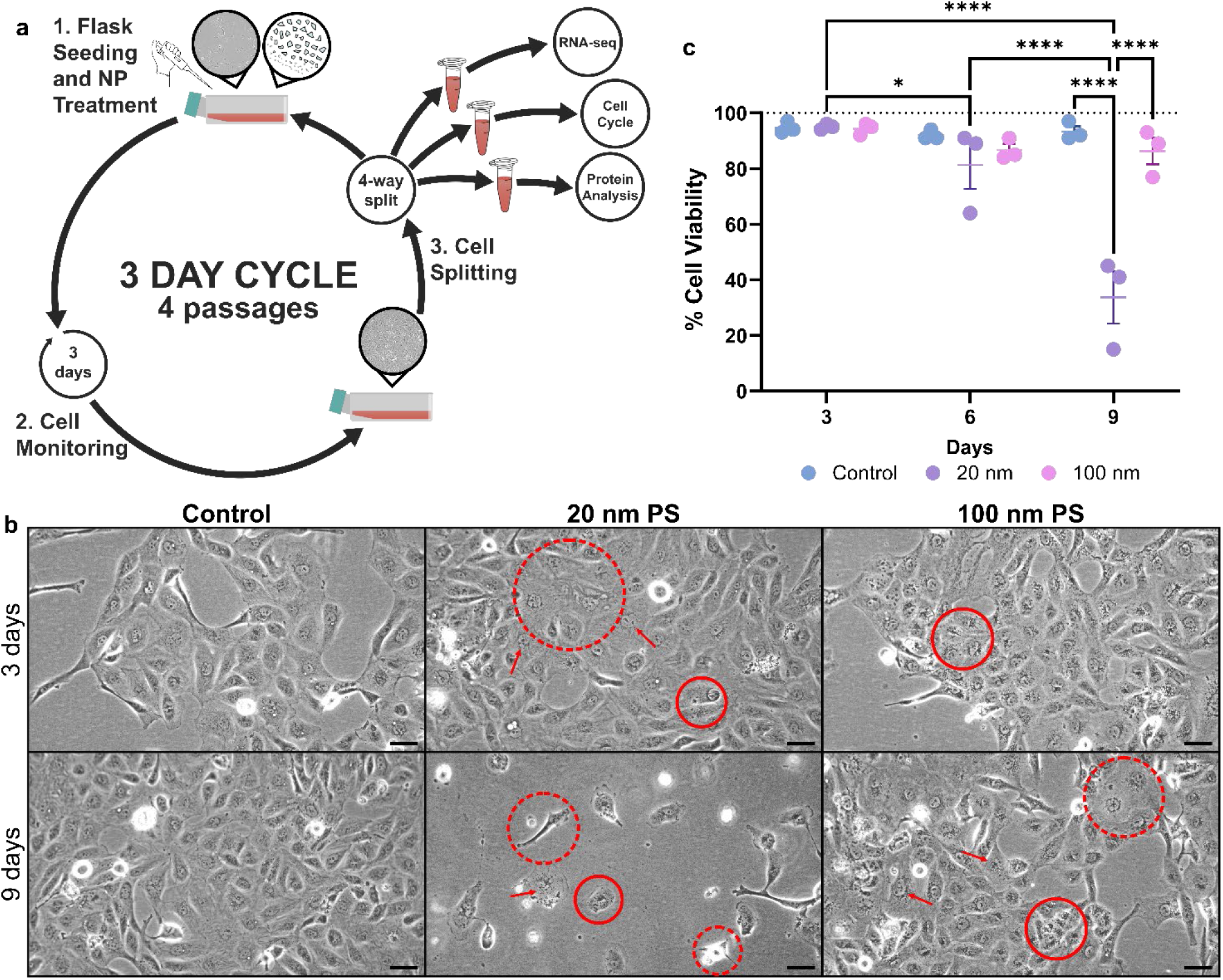
Cellular response to chronic, repeated exposure with 20 nm and 100 nm PSNPs. a) Methodology cycle for performing the long-term nanoplastic exposure study. b) Morphology of HK-2 cells at D3 and D9 of NPs exposure. Solid lines indicate regions of increased intracellular granularity. Dashed lines indicate regions of undefined or irregular cell borders. Arrows indicate nuclear irregularities. 20x magnification. Representative micrographs of intermediate timepoints are provided in Fig. S1. c) Cellular viability at D3, D6 and D9 of NPs exposure. Data in plot c is expressed as mean with standard error of the mean. N=3. P-value = ‘ns’ no significance, * ≤ 0.05, ** ≤ 0.01, *** ≤0.001, **** ≤ 0.0001.

At each timepoint, and immediately prior to cell collection and splitting, cells were imaged at the centre of each flask using a ZEISS Primovert inverted cell culture microscope with a ZEISS Axiocam 208 color/202 mono microscope camera, with a 20x objective.

### 2.3 Cell cycle analysis

Cell cycle analysis was performed to assess the effects of NPs exposure on HK-2 cells. Following each 72-hour exposure, cells (1.0 × 10^6^) were collected, washed with PBS, fixed in 80% EtOH for a minimum 24-hours, then stained with propidium iodide (50 µg/mL) in PBS containing RNase A (200 µg/mL), and 0.1% Triton X-100. Following a 30-minute incubation in the dark, samples were analysed via flow cytometry, and data was processed using FlowJo software (v10.8.1; BD Biosciences).

### 2.4 RNA-sequencing

#### 2.4.1 Sample collection and extraction

After NPs treatment, 0.1 × 10^6^ HK-2 cells were collected, pelleted and resuspended in TRIzol reagent (Invitrogen), before RNA extraction and purification using the Direct-zol RNA Microprep kit (Zymo Research) following the protocol recommended by the manufacturer. To each tube 100% EtOH was added at equal volume to TRIzol and mixed thoroughly before transfer to a spin column. All samples underwent DNase I treatment, 3 pre-wash cycles, and 1 wash cycle, before RNA elution in 20 µL of DNase/RNase-free water. Sample RNA quantity and quality was quantified via Nanodrop. A total of 26 RNA samples underwent sequencing by the South Australian Genomics Centre (SAGC). The samples were sequenced using the MGI DNBSEQ-G400, with a read length of 100 base pairs (bp), for a total 665 million total reads, or approximately 25 million reads per sample. Samples were then processed and analysed for differentially expressed genes (DEGs).

#### 2.4.2 Bioinformatics

The RNA-sequencing data was processed using the Flinders University High Powered Computer (HPC) DeepThought,^25^ using the Python package and environment manager Miniconda3 (version 22.11.1-1). Sequence trimming was performed with Cutadapt (version 5.0).^26^ Genetic index creation and alignment was performed with STAR (version 2.7.11b)^27^ using the GENCODE^28^ Human 46 GRCh38 reference genome and gene annotation. SAMtools (version 1.21)^29^ was used for sequence indexation, read summarisation was performed using the featureCounts feature within the Subread (version 2.0.8) package,^30^ and the FastQC (version 0.12.1)^31^ and MultiQC (version 1.25.2)^32^ packages were used for sequence quality checking.

All RNA-seq data analysis and figure generation was performed in RStudio (version 2024.12.1). Differential gene expression was completed using the edgeR^33^ and limma^34^ packages, with gene annotation created using AnnotationHub^35^ and the ensembldb^36^ packages. Initial gene filtration was performed with minimum thresholds of 3 counts per million and 3 samples to remove lowly expressed genes. Principal component analysis (PCA) with multidimensional scaling (MDS) was performed to assess the relationship between samples, and Surrogate Variable Analysis (SVA; version 3.54.0)^37^ was applied to remove batch effects. Significant DEGs were identified based on a log_2_ fold-change (FC) of >|1.1| and adjusted P-value of <0.05. To prepare the DEGs for gene set enrichment analysis, further filtering of the significant DEGs was done to select for highly differentiated genes with a log_2_ FC ≤-1 and ≥1. The Fast Gene Set Enrichment Analysis (fgsea) package,^38^ alongside the Molecular Signatures Database (MSigDB)^39^ package was then used to perform Hallmark pathway analysis on the highly differentiated DEGs.

#### 2.4.3 RNA determination by qRT-PCR

Measurement of RNA abundance was performed for specific genes of interest which had shown regulation in the RNA sequencing studies. Preparation of the cDNA was performed using the Superscript IV VILO Master Mix with ezDNase ezyme (Thermo Fisher Scientific), following the protocol recommended by the manufacturer. All samples were prepared to a final cDNA concentration of 5 ng/µL. Analysis of the cDNA was performed via qRT-PCR using the TaqMan Fast Advanced Master Mix (Thermo Fisher Scientific) and run on the Rotor-gene Q (Qiagen). For each primer (Table S2), each individual sample was run in triplicate. The analysis threshold was set to 0.1 for all runs. Results were calculated and expressed as FC of the delta cycle threshold (FC ΔΔCT) according to Livak and Schmittgen.^40^

#### 2.4.4 Western blotting

Protein samples were extracted from whole cell pellets using 100 µL of urea/SDS buffer (6.7 M urea, 10 mM Tris-Cl (pH 6.8), 1 mM dithiothreitol, 10% glycerol, and 1% SDS)^41^ supplemented with Halt Protease and Phosphatase Inhibitor Cocktail (100X) (ThermoFisher Scientific). Samples were then sheared by repeated passage through a 29-gauge insulin syringe, before undergoing 15 minutes of sonication in a cold-water bath. Quantification of the total protein was performed via BCA protein assay, using the Pierce BCA Protein Assay Kit (ThermoFisher Scientific), then colorimetric analysis at 562 nm using the SpectraMax iD5 Multi-Mode Microplate Reader (Molecular Devices). Final protein concentrations were determined via quadratic curve. Protein samples (20 µg) were mixed 3:1 ratio with 4x Laemmli sample buffer containing 100 mM DTT, then loaded into AnyKd Mini-PROTEAN TGX Precast Protein Gels (Bio-Rad), alongside the Chameleon Duo Pre-Stained Protein Ladder (8-260 kDa; LICORbio). The gel was run via SDS-PAGE at 100 V for 60 minutes then transferred to a PVDF membrane with a Trans-Blot Turbo Mini Transfer Pack (Bio-Rad) using the Mixed MW protocol (Bio-Rad). Blots were blocked with Odyssey Blocking Buffer (LICORbio) prior to an incubation overnight at 4°C with rabbit polyclonal angiopoietin like 4 antibody (ANGPTL4) (1:500; 18374-1-AP, Proteintech) and mouse monoclonal β-Tubulin (1:1000; D3U1W, Cell Signalling Technology) antibodies. The following day blots were washed (3 times in 1X PBS containing 0.1% Tween-20) and then incubated for 1 hour at room temperature (23°C) with anti-rabbit 800IR, and anti-mouse 680IR secondary antibodies. All blots were imaged on the Odyssey CLx Imager (LICORbio) and analysed using Image Studio (Ver. 6.1; LICORbio).

### 2.5 Statistical analysis

All experiments and analyses were run with 3 biological replicates where possible. Due to insufficient cell growth and low cell counts at D9, the 20 nm treatment group was not included in the cell cycle or protein validation analyses, and only 2 replicates were included in the RNA-seq and qRT-PCR RNA validation experiments.

Statistical analyses of the cell counts and viability, cell cycle, RNA validation and protein quantification were performed via GraphPad Prism (v10.4.0; GraphPad Software). Significance was determined with the two-way analysis of variance (ANOVA) with multiple comparisons Uncorrected Fisher’s least-significant difference test. Protein quantification was determined via quadratic curve with nonlinear regression.

## 3. Results

### 3.1 Nanoplastic characterisation

The NPs used in this study were previously characterised by Fourier transformed infrared spectroscopy (FTIR) and dynamic light scattering (DLS), which confirmed both the 20 nm and 100 nm PS NPs matched the expected spectra for PS, with hydrodynamic diameters of 48.0 nm and 131.8 nm, respectively.^23^

### 3.2 Cell morphology and viability assessment

HK-2 cells were co-cultured alongside 20 nm and 100 nm PS NPs in 3-day cycles for 4 passages to investigate the long-term effects of continuous, repeated exposure (Fig. 1a). After 12 days of NPs exposure, visual morphological differences between the control and treated cells were identified, predominantly in terms of cell granularity, and overall cell shape (Fig. 1b & S1). These changes were identifiable after the first 72-hours (D3) of NPs exposure, where abnormalities in the shape of the cell and the appearance of the nucleus had begun to appear in cells exposed to both sizes of NPs, along with increased intracellular granularity in the 100 nm group (Fig. 1b & S1). After 9-days of continued exposure, the 100 nm treatment group had a visible increase in granularity within the cytoplasm, with clear localisation around the nucleus, while the 20 nm group experienced significant reduction in the number of viable cells (Fig. 1b & c). A consistent reduction in the total number of live cells can be seen with repeated exposure to 20 nm PS NPs, reducing to 81.4% viability at day 6 (D6), and further to 33.9% viability at day 9 (D9) (Fig. 1c & S2a). This corresponds to an overall 61.1% reduction in cell viability by day 9. The 100 nm PS NP treatment group saw stable numbers of live cells at each time point, though the proportion of dead cells increased over time (Fig. 1c & S2b). At day 12, a total 0.85 × 10^6^ cells were seen in the 20 nm group (16% compared to control total), with only 0.17 × 10^6^ (19.9%) of these cells being viable (Fig. S2). The experiment was terminated after 12-days of exposure as a result.

### 3.3 RNA-sequencing and differential gene expression analysis

Following extraction and sequencing of the RNA from the PS NPs treated cells, differential gene expression analysis was undertaken to probe for timepoint associated DEGs over the 9-days of continuous NP exposure. Initial gene filtration of the sample data detected a total 12,219 genes present across the samples (Fig. S3), and library quality checking of the data identified consistent sample distribution and alignment (Fig. S4). Upon performing a MDS PCA with SVA correction, the samples were able to be separated into exposure time (PC1), and treatment (PC2) related effects (Fig. 2a). The control samples stay relatively grouped together on the PC2 axis as the timepoint increases, while both the 20 nm and 100 nm NPs treatments separate as the exposure time increases (Fig. 2a). Separation of the 20 nm and 100 nm NPs treatment groups compared to the control indicates differentiation between the effects of the two sizes (Fig. 2a). Further analysis of the highly expressed gene pool was used to identify significant DEGs (defined as adjusted P-value of <0.05). In Figure 2b-i significant DEGs are displayed in red when overexpressed and blue when under-expressed compared to the untreated control group. This analysis indicated that the 20 nm NP treatment induced the greatest change in gene expression, with 1,548 DEGs at D6 and 4,431 at D9 compared to the untreated control. In contrast, the 100 nm treatment produced only a limited transcriptional response, with 35 DEGs identified at D9 only (Fig. 2b). In the 20 nm treatment group, between the D6 and D9 timepoints, an overlap of 1,302 DEGs was observed, representing 84% of the D6 DEGs (Fig. 2c). Of the 35 DEGs identified in the 100 nm D9 group, 26 DEGs (74%) were also present in the D6 and D9 20 nm groups, with 13 DEGs which were common to all 3 DEG comparisons (Fig. 2c). Between the two 20 nm timepoints, and the 100 nm treatment group, a greater proportion of overlap was present with downregulated DEGs (695) compared to those that were upregulated (616) (Fig. S5).

**Fig. 2.**
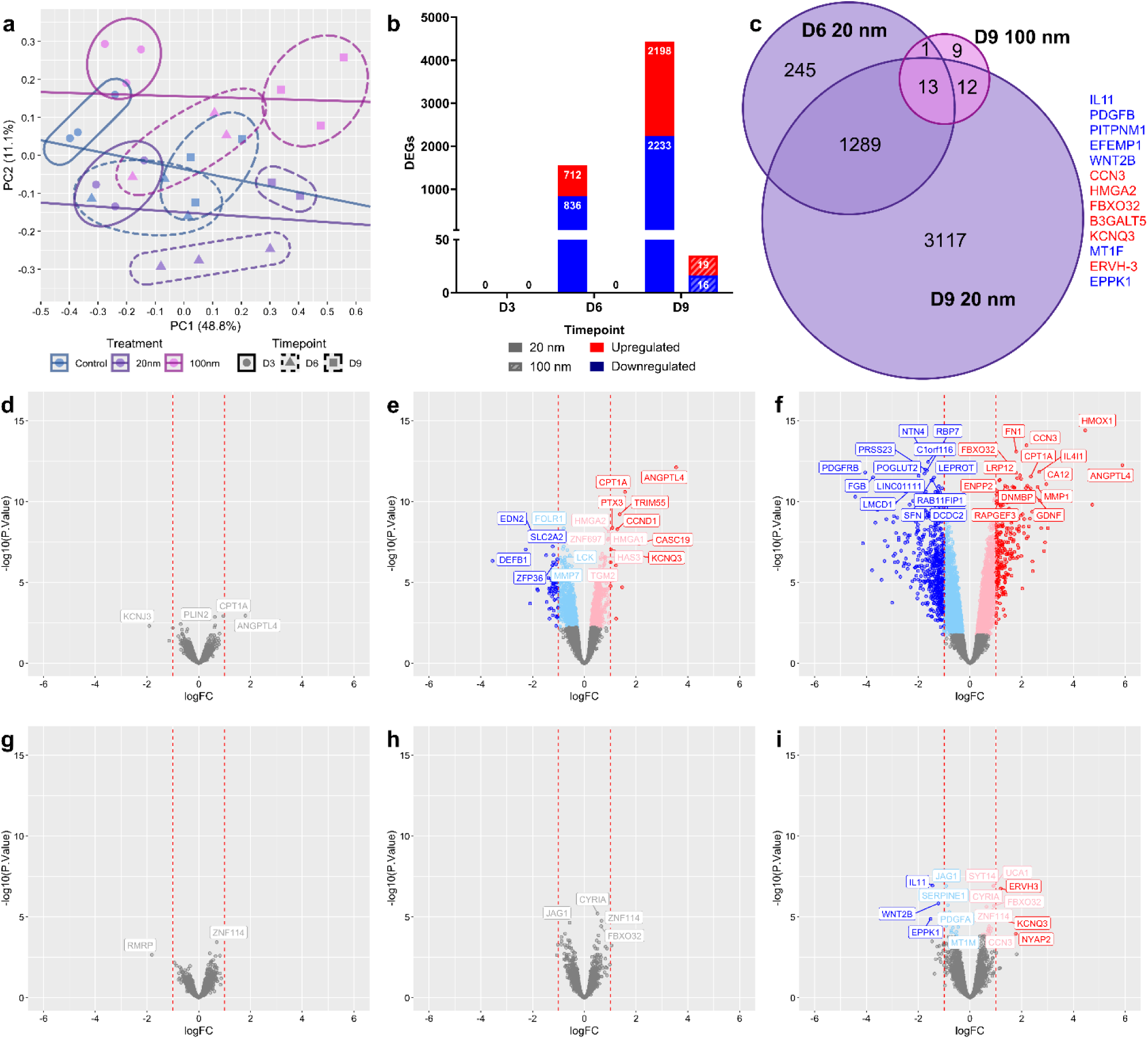
RNA-sequencing analysis of differentially expressed genes (DEGs) following long-term exposure to 20 nm and 100 nm PS NPs. a) Multidimensional Scaling (MDS) plot with Surrogate Variable Analysis (SVA) correction. b) DEG distribution at D3, D6 and D9 timepoints. c) Venn diagram showing DEG overlap between the D6 20 nm, D9 20 nm and D9 100 nm treatment groups and timepoints, with the 13 DEGs expressed in all 3 groups (listed). Red text indicates upregulated DEGs, while blue text indicates downregulated DEGs. Volcano plots of the top 30 DEGs from d) D3, e) D6 and f) D9 of the 20 nm group, and g) D3, h) D6 and i) D9 of the 100 nm treatment group. All data is expressed in comparison to the control group at each timepoint. The dashed red lines indicate the additional ≤-1 and ≥1 log_2_ FC filtering cutoff. Significant DEGs retained by this filtering are indicated as the outer blue and red points, while the light blue and pink points indicate DEGs below the log_2_ FC cutoff.

Further filtering of the significant DEGs was performed to identify those with a log_2_ FC ≤-1 or ≥1, corresponding to genes that are highly differentiated from the control (Fig 2d-i). In the 20 nm group, 74 and 760 DEGs were highly differentiated at the respective D6 (fig. 2e) and D9 (fig. 2f) timepoints, with the majority of these highly differentiated DEGs being downregulated, while only 6 DEGs were highly differentiated in the D9 100 nm group (fig. 2i). In the 20 nm group, a number of the most highly differentiated DEGs identified at the D6 timepoint were further dysregulated at the D9 timepoint, namely heme oxygenase 1 (*HMOX1*) which increased from 1.22 to 4.46 log_2_ FC (3.24 log_2_ FC difference), and *ANGPTL4* which increased from 3.55 to 5.89 log_2_ FC (2.34 log_2_ FC difference). The topmost upregulated and downregulated DEGs for the D6 and D9 20 nm treatment group, and the D9 100 nm treatment group are listed in Tables S3-S8.

An exploration of effects on cell cycle was also undertaken by examining the RNA-seq data for variation of expression of genes with characteristic cell cycle phase specific expression. The gene list collected by Hsiao et al.^42^ was used to select and filter cell cycle genes and the associated phases. This analysis identified, in the 20 nm treatment group only, a total 143 individual significant DEGs associated across all cell cycle phases of the D6 and D9 timepoints (Fig. 3a & b). Distinct trends were identified towards the M phase and M/G1 phase transition with 8 and 14 respective DEGs upregulated at D6, increasing to 28 and 25 respective DEGs at D9 (Fig. 3a). This trend saw a total 19 DEGs increasing in expression between 1-fold to 3-fold from D6 to D9. Comparatively, downregulation of the DEGs in the S and G2 phases was identified at D6 (5 DEGs – S phase; 7 DEGs – G2 phase), with both phases each increasing to 17 DEGs, and 10 genes decreasing in expression between 1.6-fold to 3.1-fold, from D6 to D9 (Fig. 3b).

**Fig. 3.**
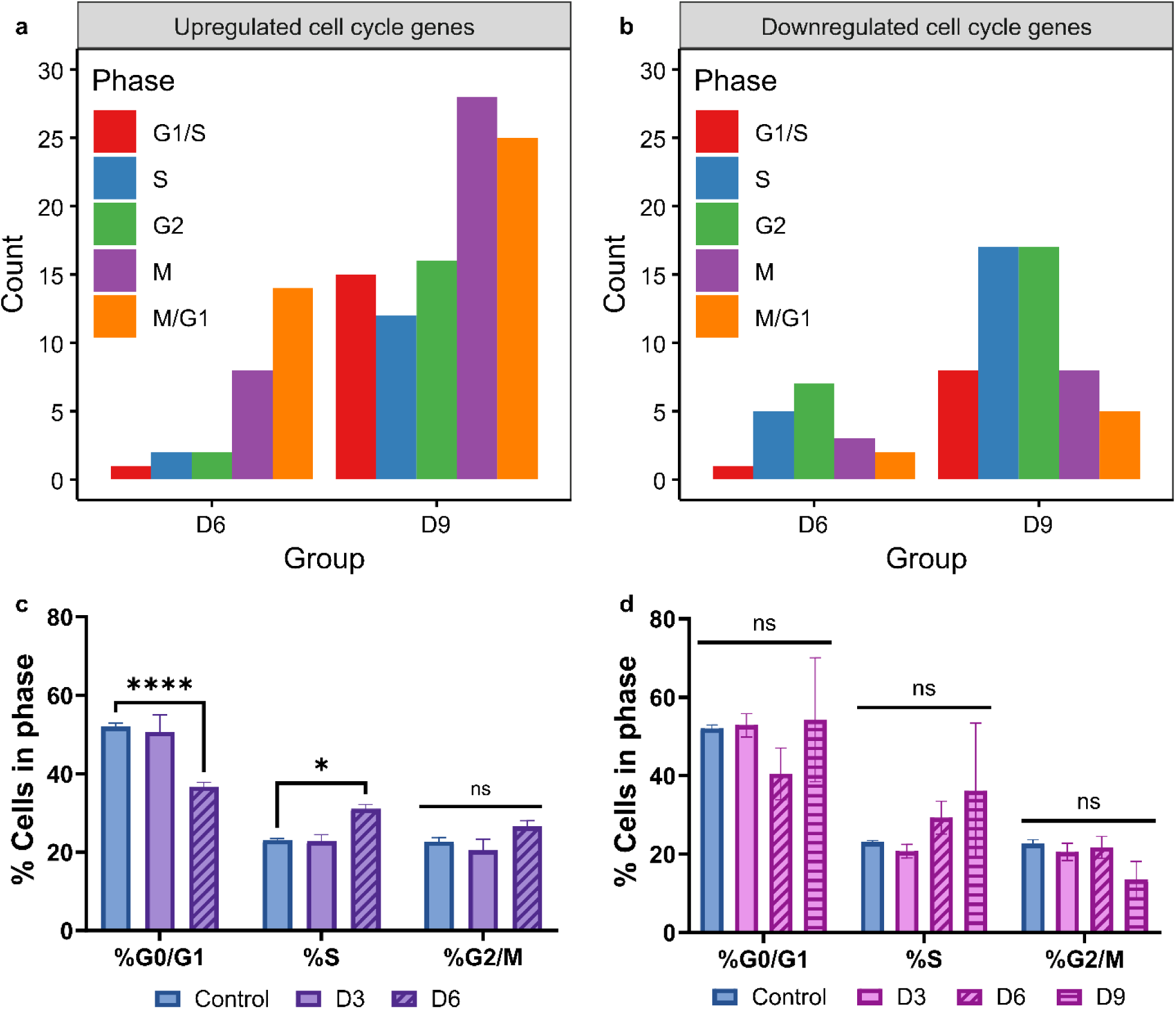
Cell cycle analysis of HK-2 cells continuously exposed to 20 nm and 100 nm PS NPs for 9 days. Comparison of significant (Adjusted P-value ≤ 0.05) a) upregulated and b) downregulated cell cycle genes in the 20 nm group following RNA-sequencing analysis. c) Flow cytometry analysis of cell phase proportions of 20 nm PS NPs treated cells at D3 and D6. Day 9 samples not included due to significant sample loss. d) Flow cytometry analysis of cell phase proportions of 100 nm PS NPs treated cells at D3, D6 and D9. Data in plots c-d are expressed as mean with standard error of the mean. N=3. P-value = ‘ns’ no significance, * ≤ 0.05, ** ≤ 0.01, *** ≤0.001, **** ≤ 0.0001.

Following repeated exposure to the PS NPs, cell cycle analysis was performed via flow cytometry to assess for potential effects on cell growth and proliferation (Fig. 3c & d). After D6 of exposure to the 20 nm NPs, a significant decrease was observed in the percentage of cells in the G0/G1 phase, coupled with a significant increase in the percentage of cells in the S phase (Fig. 3c). Analysis of the D9 timepoint in the 20 nm group was unable to be performed due insufficient cell numbers as a direct result of the reduced viability (Fig. 2b & 3c). No significant changes to cell cycle were observed up to D9 of exposure to 100 nm PS NPs, although large variation in the standard error of the D9 group was observed (Fig. 3d).

In order to explore the effects of NP exposure on biological functions, hallmark pathway analysis of the DEGs from the 20 nm (D6 and D9) groups was undertaken (Fig. S6). A total 33 individual pathways were identified with strong correlation to the DEGs at the corresponding timepoints, with 6 pathways common to both timepoints but only 4 significant pathways, all in the D9 group (Tables S9 & S10; Fig. 4a & S7). The pathway with the strongest significance was the *KRAS SIGNALING UP* pathway, showing a trending increase between D6 and D9, with a normalised enrichment score (NES) of 2.4 and an adjusted P-value of 0.003 (Fig. 4a). A total of 13 leading genes are associated with this pathway at D9, with 3 genes overlapping between the two timepoints – *ANGPTL4*, transmembrane protein 158 (*TMEM158*) and colony stimulating factor 2 (*CSF2*) (Tables S9 &S10; Fig. 4b). In the D9 20 nm treatment group, significant upregulation of the *IL2 STAT5 SIGNALLING*, *G2M CHECKPOINT*, and *FATTY ACID METABOLISM* pathways was also identified, each with an adjusted P-value of 0.04 (Fig. 4a). None of these pathways were identified in the D6 20 nm group (Fig. S6a), and no pathways were identified in the 100 nm (D9) group.

**Fig. 4.**
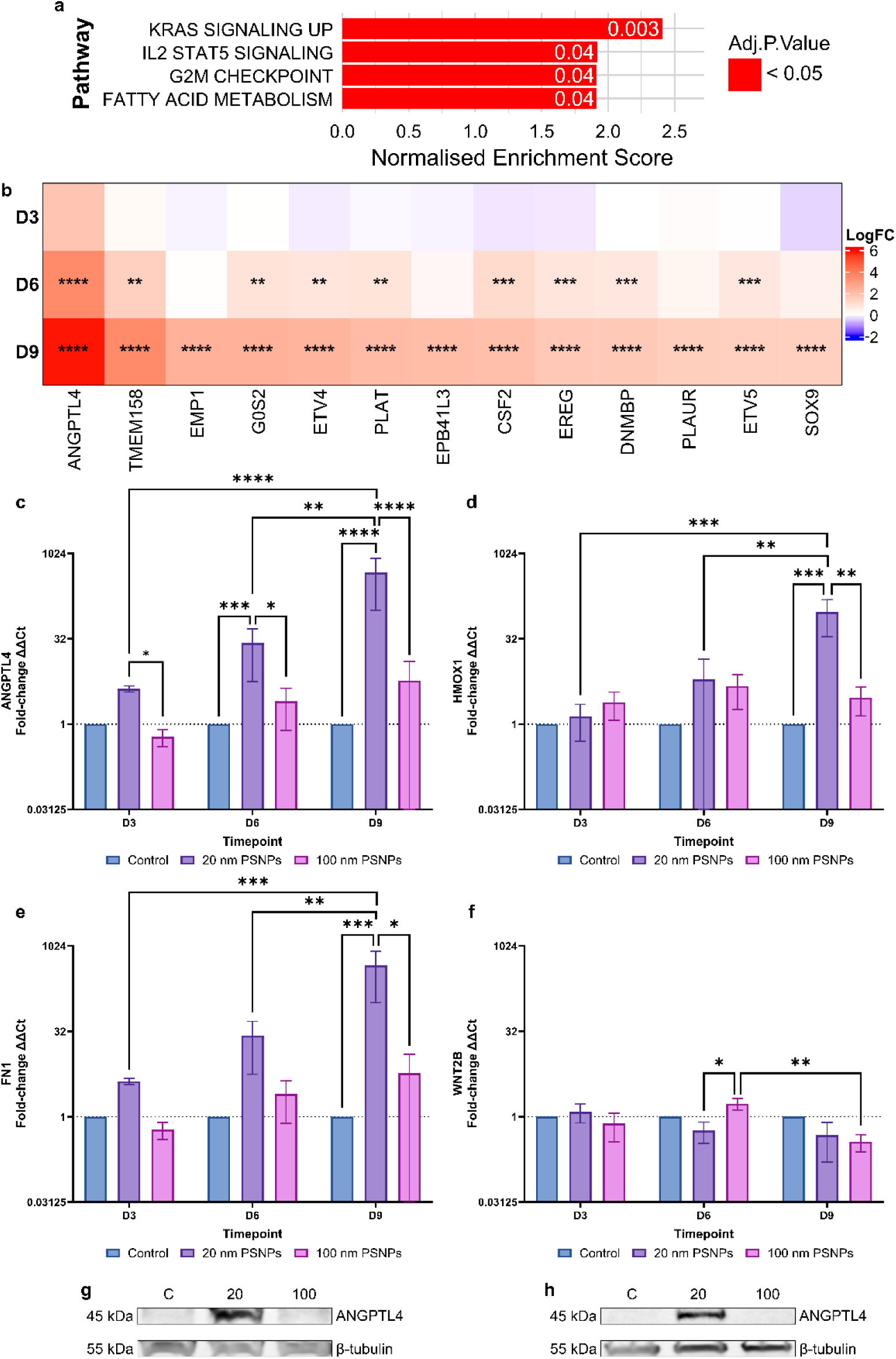
Pathway analysis, and mRNA and protein induction following PS NPs exposure. a) Top hallmark pathways following exposure to 20 nm PSNPs at day 9. DEGs were filtered to select for those with a log2 FC of ≥1 or ≤-1. b) Heatmap of the 13 DEGs identified as the leading genes in the KRAS Signalling Up pathway, showing the log2 FC expression at each timepoint. Red indicates upregulation, blue indicates downregulation. Significance is adjusted P-value. ** ≤0.01, *** ≤0.001, **** ≤0.0001. Data is associated with Tables S9 & S10. RNA validation of key DEGs, c) ANGPTL4, d) HMOX1, e) FN1, and f) WNT2B via qRT-PCR analysis. N=3. P-value = * ≤ 0.05, ** ≤ 0.01, *** ≤0.001, **** ≤ 0.0001. Data expressed as fold-change of the ΔΔ cycle threshold (CT). Protein expression of ANGPTL4 was validated by treating HK-2 cells with g) 100 µg/mL (N=3) or h) 1,000 µg/mL (N=2) of 20 nm and 100 nm PS NPs for 24-hours before collection. Whole-cell lysates were separated by SDS–PAGE and transferred onto PVDF membranes. Membranes were probed for ANGPTL4 (∼45 kDa) and β-tubulin (∼55 kDa), which was used as the loading control. Protein bands were visualised using the LI-COR Odyssey imaging system.

### 3.4 RNA and protein expression of selected genes

To validate the mRNA expression identified through the RNA-sequencing analysis, qRT-PCR was performed of some of the most highly differentiated genes across the D6 and D9 timepoints of both treatment groups. *ANGPTL4* and *HMOX1* were selected due to their significant dysregulation in the 20 nm treatment group at both the D6 and D9 timepoints (Tables S3 & S5). Fibronectin 1 (*FN1*) was selected due to its significant induction at D9 in the 20 nm treatment group (Table S5), and Wnt family member 2B (*WNT2B*) was selected for its significant downregulation across both the 20 nm and 100 nm treatment groups (Tables S6 & S8). All four DEGs additionally have associations with kidney conditions such as renal fibrosis and acute kidney injury (AKI).^15, 43–45^

Of the 4 genes, *ANGPTL4*, *HMOX1* and *FN1* all confirmed increased expression, as had been indicated with the RNA-seq analysis (Fig. 4c-e). RNA-sequencing confirmed *ANGPTL4* as the DEG with the greatest overall upregulation at both the D6 (3.55 log_2_ FC) and D9 (5.89 log_2_ FC) timepoints in the 20 nm treatment group, which aligned with the qRT-PCR validation values of 26.8 FC ΔΔCt at D6 and 467.9 FC ΔΔCt at D9, equivalent to a log_2_ FC of 4.7 and 8.9 when compared to the respective timepoint’s control group (Fig. 4c). RNA expression of *ANGPTL4* was shown to increase in a time-dependent manner as exposure to the 20 nm NPs continued, with a 6-fold change between D3 and D6, and a further 17-fold change between D6 and D9 (Fig. 4c). Expression of *HMOX1* and *FN1* in the 20 nm treatment group was also significantly increased in comparison to the control at D9, with respective 95-fold and 16-fold increases in expression (Fig. 4d & e). The only significant changes in expression validated via qRT-PCR for the 100 nm NPs treatment group was *WNT2B*, which expressed a significant reduction in expression from D6 to D9 (21.4% decrease), which was also identified in this treatment group in the RNA-seq data (Fig. 4f, Table S8). In both the *ANGPTL4* and *FN1* validations, a trend of time-dependent, though non-significant, expression increases can be seen (Fig. 4c & e).

To confirm the protein expression of ANGPTL4 following NPs exposure, HK-2 cells were exposed for 24-hours to 100 µg/mL and a higher 1 mg/mL concentration of PS NPs. Following exposure, ANGPTL4 expression could be seen in the 20 nm group at both tested concentrations (Fig. 4g & h). No visible expression was identified following exposure with the 100 nm PS NPs (Fig. 4g & h).

## 4. Discussion

Increasing concern for the health implications of daily exposure to NPs has led to rapid progression in NPs toxicity research over the last five years.^46, 47^ With the known prevalence and accumulation of MPs and NPs within environmental and biological matrices,^1, 48^ studies assessing the health impacts of continuous, frequent exposure should be the primary focus. Despite these advances, current research is predominantly focused on assessing the short-term, acute exposure effects, leaving the biological impacts of chronic, repeated NPs exposure both understudied and poorly understood.^49^ To address this research gap, we developed a long-term *in vitro* exposure methodology to investigate how continuous, repeated exposure of NPs alters the morphology, viability and transcriptional regulation of HK-2 cells, whilst also assessing the influence of NPs size on these responses.

Through continuous co-culture of HK-2 cells with PS NPs, in as little as 3 days, identifiable morphological changes were seen, particularly involving the 20 nm NPs (Fig. 1c). At D3 both NPs treatment groups expressed increased cell granularity, localised close to the nucleus. This feature was more prominent in the 100 nm group and followed through to D9, at which point the 20 nm group had decreased to 12% of the total number of viable cells (Fig. 1c & S1a). A recent study by Kim et al.^49^ similarly investigated long-term NPs exposure *in vitro*, with 20 nm, 50 nm and 100 nm PS NPs, though their study utilised the human colorectal cancer (HCT116) cell line. In their study, the HTC116 cells were co-cultured with the PS NPs, passaging the cells and replenishing the media every 2-3 days, for a total 2 months and 21 total passages.^49^ Throughout the treatment period, morphological changes were identified around passages 4-5 following exposure to the 20 nm PS NPs at 10 µg/mL and 20 µg/mL, with the higher concentration showing the greatest effect; no significant effect was seen with the larger 50 nm and 100 nm NPs.^49^ The effects seen by Kim et al.^49^ show similar trends to the results seen in our study, where exposure to the 20 nm NPs significantly impact morphology and viability, and minimal impact is caused by the larger 100 nm NPs (Fig 1c). It’s worth noting that the concentrations used by Kim et al.^49^ were 5- and 10-times lower than the 100 µg/mL concentration tested in our study, but it was not made clear at what point of the cell culture protocol the NP treatments were added to the cells after passaging (during seeding or post-monolayer formation).

The study by Zhu et al.^20^ provides a more relevant comparison to our study as it assessed the impacts of 20 nm and 50 nm PS NPs on HK-2 cells, at concentrations of 50 µg/mL and 100 µg/mL, albeit the focus of this study was limited to a short-term, single dosage period of 12 hours. Comparatively, this study identified significant reductions in viability at 100 µg/mL, with the greatest effect also seen in the smaller 20 nm NPs treatment group, reducing to approximately 60% viability, where exposure to the 50 nm NPs dropped to approximately 90% viability.^20^ In our study, it was only at D6 that we saw a significant reduction in viability in the 20 nm group, dropping only to 81% viability, and it was not until D9 where we saw the viability further reduce to 34% (Fig 1b). Given the increased cell response of the 20 nm PS NPs at 100 µg/mL, Zhu et al.^20^ proceeded with RNA-sequencing analysis of the samples, identifying a total 1,379 DEGs (683 up, 696 down). At the D6 timepoint, our 20 nm PS NPs treated cells expressed a similar proportion of DEGs with 712 upregulated, and 836 downregulated (Fig. 2b & e).

Kim et al.^49^ also reported a short-term cell cycle analysis of the 20 nm PS NPs treatment group, with concentrations up to 50 µg/mL, ran via flow cytometry, though no significant changes between phases or concentrations were identified.^49^ Comparatively, our previous short-term exposure study identified dose-dependent shifts from 0.2 µg/mL (7.8×10^9^ particles/mL) to 200 µg/mL (7.8×10^12^ particles/mL – equivalent to this study’s concentration of 100 µg/mL; Table S1) in cell cycle for the 20 nm PS NPs after 24 hours, where the proportion of cells in the G0/G1 phase was reduced, and the S phase proportion was increased.^23^ Interestingly in the long-term study presented here, no significant cell cycle effects were seen with the 100 µg/mL dose at D3, but a similar pattern to the short-term study was identified after D6 of exposure, indicating ongoing impact to the cell cycle (Fig. 3a). This effect is likely to be attributed to the reduced viability at the D6 timepoint, and the significant (Adjusted P-value 0.04) upregulation of the G2M checkpoint hallmark pathway (Fig. 1b & 4a). The spikes of dysregulation identified from the RNA-seq cell cycle analysis, specifically down regulation of genes involved with cycling through S to G2 phase and concomitant up regulation of genes involved with the cell cycle phase from M to G1 seem to align with the S-phase arrest and decrease in the proportion of cells in the G0/G1 phase noted in the flow cytometry analysis (Fig. 4a). While the individual, per cell, cell cycle phases are impossible to distinguish due to the nature of bulk RNA-seq data, the alignment of the flow cytometry cell cycle data with the RNA-seq output likely reflects a true population-level effect resulting from sustained NPs exposure. Another potential source of cell cycle dysregulation is through the *KRAS* signalling pathway. Activation of the *KRAS* signalling pathway is responsible for downstream regulation of cell proliferation, differentiation and apoptosis, depending on the activated pathway,^50^ though most commonly, *KRAS* activation is associated with oncogenic mutations, often indicating cancer severity and tumour formation.^50, 51^ Importantly, activation of *KRAS* has previously been seen through the induction of oxidative stress and increased reactive oxygen species (ROS) production, processes which are frequently associated with exposure to MPs and NPs,^52–54^ and also play a key role in CKD progression.^14, 55, 56^

The most highly dysregulated gene identified within the *KRAS* signalling pathway, whilst also being the most upregulated DEG at both D6 and D9 in the 20 nm group, was *ANGPTL4* (Table S3 & S5). ANGPTL4 is a glycoprotein responsible for regulating fatty acid metabolism through inhibition of lipoprotein lipase (LPL), as well as involvement in regulating angiogenesis, inflammation, and oxidative stress.^57–60^ While ANGPTL4 is expressed primarily in the liver and adipose tissue,^61^ its overexpression has been identified as a key regulator for a number of kidney conditions including CKD, diabetic kidney disease (DKD) and AKI.^58, 62–64^ Overexpression of ANGPTL4 in CKD, DKD and AKI has been linked to the regulation of fibrosis,^62^ and the development and progression of renal interstitial fibrosis (RIF).^15, 65^ Knockout of ANGPTL4 in diabetic mice causes suppression of renal fibrosis development.^63^ Induction of ANGPTL4 protein expression in addition to mRNA induction was identified in the 20 nm PS NPs treatment group after 24-hours of exposure at 100 µg/mL, with no expression seen in the control or 100 nm PS NPs treatment groups (Fig. 4g). Marked induction was additionally seen following an increase in NPs concentration to 1 mg/mL (Fig. 4h).

Furthermore, RNA validation of *ANGPTL4* confirmed the increased expression identified through RNA-seq analysis, along with *HMOX1* and *FN1* (Fig. b-d), all of which have reported roles in the progression of fibrosis in CKD, AKI and DKD.^43, 63, 64^ The induction of kidney fibrosis following long-term exposure to MPs was recently identified in a study by Hong et al..^66^ In this study male BALB/c mice were provided with 10 mg/L of 5 µm PS MPs (unmodified or amine (NH_2_)-modified) in drinking water for a total 6 months, after which significant regions of fibrosis were identifiable within the kidney tissue, with the more severe effect seen in the NH_2_-modified MPs.^66^ The mechanism underlying *ANGPTL4* induction (and the other regulated genes) following NPs exposure remains to be established, though several plausible pathways exist. *ANGPTL4* is a well-characterised transcriptional target of the peroxisome proliferator-activated receptor (PPAR) family, particularly PPAR-γ, and is also induced downstream of hypoxia-inducible factor (HIF) signalling and cellular oxidative stress,^15, 57, 58, 61^ both of which have been implicated in NP- and MP-induced cellular responses.^67, 68^ Given the concurrent activation of the KRAS pathway and suggestion of oxidative stress observed in this and other studies, it is plausible that sustained NP exposure induces *ANGPTL4* via ROS-mediated activation of PPAR and/or HIF signalling, though direct testing of these pathways (e.g., through pathway-specific inhibition) will be required to confirm this mechanism. The prominent induction of ANGPTL4 is also noteworthy because ANGPTL4 regulates fatty acid metabolism through inhibition of LPL. This lipid-related function is relevant in the context of current debate around mass-spectrometry-based detection of nanoplastics in biological tissues,^68–70^ where endogenous lipids can generate overlapping pyrolysis products and complicate polymer assignment.^71^ Although this analytical limitation is distinct from the biological response measured here, the strong dysregulation of a lipid-regulatory gene raises the possibility that nanoplastic exposure may itself perturb fatty acid metabolism, adding a further layer of complexity when interpreting associations between plastic burdens and lipid-rich tissue compartments.

Studies extending beyond the commonly used 24–48-hour NPs exposure window remain limited, particularly those involving exposure during cell seeding, when cells are still in suspension within the media and are more likely to encounter NPs than after attachment and monolayer formation. This distinction is important because, as shown in a study by Moore et al.,^72^ differences in the administration method of nanoparticles influences the particle-cell interaction. In this study, 116 nm gold nanoparticles, 100 nm silica nanoparticles, and 22 nm super-paramagnetic iron oxide nanoparticles (SPIONs) were administered to the murine J774A.1 macrophage cell line at 70 µg/mL as either a single concentrated dose into the existing cell media without mixing, a single concentrated dose into the existing cell media which was mixed via pipetting, or pre-mixed into the cell media prior to exposure.^72^ After 24-hours of exposure, both the gold and silica nanoparticles had the greatest uptake after administration of the concentrated unmixed dose while the pre-mixed dose displayed the least uptake.^72^ Comparatively, the smaller SPIONs saw limited uptake within the J774A.1 cells, and no difference between the dosage method, indicating a size-dependent difference in particle uptake.^72^ These findings highlight that NP administration method can substantially influence particle–cell interactions and uptake. Therefore, greater clarity is needed between studies, on when NPs are administered following cell seeding, as well as how they are administered.

Overall, the size of NPs appears to be directly associated with the toxicity and response of HK-2 cells, with larger particles having a lesser or delayed impact, and the smaller particles displaying greater toxicity and genetic dysregulation. These differences are evident in comparison of cell viability, cell cycle regulation and differential gene expression, as the exposure time increases (Fig. 1, 2, 3c & d). The size-dependence observed in this study is consistent with the broader NP toxicology literature and may reflects differences in cellular uptake or in toxicity mechanisms. Smaller NPs have a substantially greater surface-area-to-volume ratio, increasing the proportion of surface-exposed carboxyl groups available for interaction with the cell membrane and extracellular proteins,^73^ and favouring uptake via clathrin- and caveolae-mediated endocytosis.^9^ Once internalised, smaller NPs might be more able to escape or bypass endolysosomal compartments and accumulate within the cytoplasm and perinuclear region, whereas larger particles may be sequestered within endolysosomal compartments where their bioreactivity is comparatively limited. This potential for the 20 nm NPs to have a greater degree of intracellular accumulation and membrane interaction offers a plausible explanation for the substantially greater cytotoxic, cell-cycle and transcriptional disruption observed relative to the 100 nm NPs across all timepoints in this study.

Future work would benefit from further repeats of the experiment with NPs of varying polymers, shapes and sizes, including mixed polymer samples, and lower, more environmentally representative concentrations of NPs to improve understanding of repeat exposure and accumulation within the renal system. Single cell RNA-sequencing would also be of benefit for future analysis, as this would allow for probing of cellular changes at the individual level. Further investigation into the activation of *ANGPTL4* to determine the mechanism of induction following NPs exposure, particularly early induction, could designate *ANGPTL4* as a potential marker for early NPs stress. The disparity in cell cycle effects between the 24-hour exposure time in our short-term study^23^ and the 72-hours of exposure to NPs in this study may indicate re-regulation of the cell cycle between these timepoints. Completion of a study of exposure to NPs with intermediate assessment timepoints between 24- and 72-hours would benefit for determining the onset of cell-cycle dysregulation.

This study has several limitations. Firstly, experiments were conducted in a single, immortalised cell line (HK-2); while HK-2 cells are a widely used and well-characterised model of the proximal tubule, findings may not fully extend to primary human proximal tubule cells, other nephron segments, or the kidney *in vivo*, where NP handling and cellular responses may differ. Secondly, carboxyl-modified PS NPs were tested at a single surface chemistry in two discrete sizes (20 nm and 100 nm). Environmental NPs are considerably more heterogeneous; comprising of multiple polymer types, irregular particle shapes, weathered surfaces, and likely adsorption of environmental pollutants, all of which may contribute to different uptake and toxicity profiles. Thirdly, the treatment concentration (100 µg/mL) exceeds current estimates of environmental and dietary NPs exposure, and, together with the constant presence of NPs from the time of seeding, these factors may accentuate cellular responses relative to lower-level exposures encountered *in vivo*. Finally, this study is limited to *in vitro* endpoints; while the upregulation of *ANGPTL4*, *HMOX1* and *FN1* is consistent with pathways implicated in renal fibrosis, no functional fibrotic outcomes (e.g., collagen deposition, epithelial-to-mesenchymal transition markers, or TGF-β signalling) were assessed, and *in vivo* confirmation will be required to establish whether chronic NPs exposure does contribute to fibrotic kidney disease via these pathways.

## 5. Conclusion

This study established a long-term *in vitro* exposure model demonstrating that continuous NP exposure elicits significant, size-dependent cytotoxic and transcriptional effects in human proximal tubule cells. Chronic exposure to 20 nm PS NPs induced cell death and transcriptional dysregulation within 9 days, effects that were not seen with exposure to 100 nm PS NPs. While the 100 nm group showed minimal effects within 9 days, nearly three-quarters of the differentially expressed genes that were identified overlapped with those regulated in the 20 nm exposure group. This suggests a similar, less marked or delayed response to the larger particle size. In the 20 nm group, significant and progressively worsening changes to cell morphology, cell cycle progression, and gene expression were observed with increasing exposure duration, highlighting the cumulative, time-dependent nature of NP-induced cellular stress. Notably, this chronic exposure model revealed clear links to clinically relevant kidney conditions, including CKD and AKI, primarily through the marked and sustained upregulation of *ANGPTL4*, a key regulator of renal fibrosis. These findings suggest *ANGPTL4* as a promising candidate biomarker of NP-induced renal stress and underscore the importance of assessing chronic, repeated exposures, rather than short-term, single-dose models, when evaluating the renal health risks of environmental NPs.

## Acknowledgements

This work was supported by the Australian Research Council Future Fellowship Grant (FT200100301), The Flinders Foundation and the Flinders Medical Centre Renal Research Fund. The authors acknowledge the South Australian Genomics Centre (SAGC) which provided bulk mRNA-sequencing. The SAGC is supported by the National Collaborative Research Infrastructure Strategy (NCRIS) via Bioplatforms Australia and by the SAGC partner institutes. We acknowledge and thank Dr. Giles Best at the Flow Cytometry Facility and the Omics Facility at Flinders University.

