## Supplemental Information for "Chronic nanoplastic exposure causes size-dependent cytotoxicity and ANGPTL4 induction in human kidney cells"

Affiliations:

### Supplementary Information

#### Supplementary Tables

Table S1 Summary of NPs treatment concentrations and particle numbers. *Details provided by manufacturer.

| Supplier | Polymer | Manufacturer Label* (nm) | Manufacturer Reported Diameter* (nm) | Stock Concentration*  (% w/v) | Cat No. | Lot No. | Density (g/cm^3^) | Dilution  Concentration (µg/mL) | Particles/mL | Particles/Cell |
| --- | --- | --- | --- | --- | --- | --- | --- | --- | --- | --- |
| Thermo Fisher Scientific | PS | 20 | 28 | 4.1 | C37261 | 2418536 | 1.055 | 100 | 7.80x10^12^ | 7.8x10^7^ |
|  |  | 100 | 110 | 4.5 | C37485 | 2646517 |  |  | 1.36x10^11^ | 1.36x10^6^ |

Table S2 TaqMan primer details

| Primer | Assay ID |
| --- | --- |
| Angiopoietin like 4 (ANGPTL4) | Hs01101127_m1 |
| Heme Oxygenase 1 (HMOX1) | Hs01110250_m1 |
| Fibronectin 1 (FN1) | Hs01549976_m1 |
| Wnt family member 2B (WNT2B) | Hs00921614_m1 |
| Beta-2 microglobulin (B2M) (Housekeeper) | Hs00187842_m1 |

Table S3 Top 16 upregulated DEGs ranked by adjusted P-value (adj.P.Val) with a logFC of ≥1 from D6 of the 20 nm PS NPs treatment group. Associated with Fig. 2

| Gene Symbol | Description | logFC | AveExpr | t | P.Value | adj.P.Val |
| --- | --- | --- | --- | --- | --- | --- |
| ANGPTL4 | angiopoietin like 4 | 3.549387286 | 2.949363449 | 13.57138678 | 7.80E-13 | 9.53E-09 |
| CPT1A | carnitine palmitoyltransferase 1A | 1.571613571 | 1.87346502 | 11.35752261 | 2.58E-11 | 1.58E-07 |
| TRIM55 | tripartite motif containing 55 | 1.370757868 | 4.424174948 | 9.642037475 | 6.24E-10 | 2.54E-06 |
| PTX3 | pentraxin 3 | 1.085405491 | 2.822113208 | 8.680980021 | 4.33E-09 | 1.06E-05 |
| CCND1 | cyclin D1 | 1.277366835 | 5.315936696 | 8.596029943 | 5.22E-09 | 1.06E-05 |
| CASC19 | cancer susceptibility 19 | 2.108524352 | 2.587011174 | 7.663563397 | 4.24E-08 | 4.70E-05 |
| KCNQ3 | potassium voltage-gated channel subfamily Q member 3 | 1.021237388 | 3.050186402 | 7.277540314 | 9.25E-08 | 7.16E-05 |
| NT5E | 5'-nucleotidase ecto | 1.240561225 | 4.822244875 | 6.543287267 | 5.17E-07 | 0.000167743 |
| ENSG00000260877 | novel transcript | 1.504515344 | 0.367826811 | 6.549459722 | 5.22E-07 | 0.000167743 |
| FRMD3 | FERM domain containing 3 | 1.01729235 | 1.707638602 | 6.489097629 | 5.80E-07 | 0.000177057 |
| HMOX1 | heme oxygenase 1 | 1.220602979 | 6.023658857 | 6.30597104 | 9.10E-07 | 0.000231663 |
| CSF2 | colony stimulating factor 2 | 1.135130701 | 2.715434398 | 5.530356528 | 6.02E-06 | 0.000736216 |
| SHC4 | SHC adaptor protein 4 | 1.099826105 | 0.716132796 | 5.426593437 | 7.77E-06 | 0.000821566 |
| FAM222A | family with sequence similarity 222 member A | 1.012327695 | 1.39815048 | 5.096417969 | 1.76E-05 | 0.001230645 |
| TMEM158 | transmembrane protein 158 | 1.463314703 | 0.616200631 | 5.060589409 | 2.03E-05 | 0.00135886 |
| SIGLEC15 | sialic acid binding Ig like lectin 15 | 1.228262446 | 0.111214839 | 3.281639155 | 0.00181138 | 0.020704628 |

Table S4 Top 20 downregulated DEGs ranked by adjusted P-value (adj.P.Val) with a logFC of ≤-1 from D6 of the 20 nm PS NPs treatment group. Associated with Fig. 2

| Gene Symbol | Description | logFC | AveExpr | t | P.Value | adj.P.Val |
| --- | --- | --- | --- | --- | --- | --- |
| EDN2 | endothelin 2 | -1.927247077 | 2.431493168 | -8.032249247 | 1.84E-08 | 2.70E-05 |
| SLC2A2 | solute carrier family 2 member 2 | -1.222830405 | 2.274577211 | -7.460946391 | 6.17E-08 | 5.80E-05 |
| DEFB1 | defensin beta 1 | -2.264475537 | 1.3026716 | -7.319157444 | 9.38E-08 | 7.16E-05 |
| ATF3 | activating transcription factor 3 | -1.191891833 | 6.92745306 | -6.941843202 | 2.02E-07 | 0.000129213 |
| SERPINF2 | serpin family F member 2 | -1.479095188 | 3.101699263 | -6.926523331 | 2.14E-07 | 0.000129213 |
| ZFP36 | ZFP36 ring finger protein | -1.042231901 | 6.488895274 | -6.662565844 | 3.85E-07 | 0.000156773 |
| EGR3 | early growth response 3 | -3.549299756 | 0.027526904 | -6.673657304 | 4.77E-07 | 0.000166695 |
| PSG3 | pregnancy specific beta-1-glycoprotein 3 | -1.20260917 | 4.067089155 | -6.500863642 | 5.70E-07 | 0.000177057 |
| SLC17A1 | solute carrier family 17 member 1 | -1.089650824 | 2.001583946 | -6.367783344 | 7.77E-07 | 0.000215869 |
| FOS | Fos proto-oncogene, AP-1 transcription factor subunit | -1.229846254 | 8.349784067 | -6.313779317 | 8.94E-07 | 0.000231663 |
| RARRES2 | retinoic acid receptor responder 2 | -1.108657273 | 2.934009824 | -5.872012238 | 2.59E-06 | 0.00046829 |
| CNN1 | calponin 1 | -1.00157949 | 2.687602793 | -5.799035938 | 3.07E-06 | 0.000500204 |
| PIGZ | phosphatidylinositol glycan anchor biosynthesis class Z | -1.174107184 | 1.897792541 | -5.600582971 | 5.08E-06 | 0.000674671 |
| USH1C | USH1 protein network component harmonin | -1.395092325 | 1.612839552 | -5.58736085 | 5.37E-06 | 0.000690436 |
| TCP11L2 | t-complex 11 like 2 | -1.286328364 | 2.265159886 | -5.537176933 | 6.01E-06 | 0.000736216 |
| CIMAP1B | ciliary microtubule associated protein 1B | -1.136968691 | 1.227872534 | -5.496944253 | 6.54E-06 | 0.000776307 |
| LINC02532 | long intergenic non-protein coding RNA 2532 | -1.784204559 | 1.091696181 | -5.479430812 | 7.38E-06 | 0.000821566 |
| NR4A1 | nuclear receptor subfamily 4 group A member 1 | -1.618802811 | 3.768521478 | -5.453579187 | 7.70E-06 | 0.000821566 |
| ADGRA2 | adhesion G protein-coupled receptor A2 | -1.275775386 | 2.457569622 | -5.435134194 | 7.74E-06 | 0.000821566 |
| BMF | Bcl2 modifying factor | -1.14568403 | 3.173783681 | -5.407696127 | 8.18E-06 | 0.000821566 |

Table S5 Top 20 upregulated DEGs ranked by adjusted P-value (adj.P.Val) with a logFC of ≥1 from D9 of the 20 nm PS NPs treatment group. Associated with Fig. 2

| Gene Symbol | Description | logFC | AveExpr | t | P.Value | adj.P.Val |
| --- | --- | --- | --- | --- | --- | --- |
| HMOX1 | heme oxygenase 1 | 4.458842203 | 6.023658857 | 17.3983892 | 3.99E-15 | 4.88E-11 |
| CCN3 | cellular communication network factor 3 | 2.187950252 | 3.522677905 | 15.65533717 | 3.34E-14 | 2.04E-10 |
| FN1 | fibronectin 1 | 1.791629346 | 12.76883854 | 14.99222725 | 8.22E-14 | 2.51E-10 |
| ANGPTL4 | angiopoietin like 4 | 5.891465457 | 2.949363449 | 13.85460618 | 5.81E-13 | 1.18E-09 |
| IL4I1 | interleukin 4 induced 1 | 2.678481019 | 2.579685263 | 13.10830644 | 1.50E-12 | 1.89E-09 |
| FBXO32 | F-box protein 32 | 1.78350738 | 5.845445563 | 12.98353948 | 1.71E-12 | 1.89E-09 |
| ERVH-3 | endogenous retrovirus group H member 3 | 1.934875797 | 4.577821091 | 12.76763616 | 2.44E-12 | 2.29E-09 |
| CPT1A | carnitine palmitoyltransferase 1A | 2.3532437 | 1.87346502 | 12.69426269 | 2.84E-12 | 2.31E-09 |
| ZNF697 | zinc finger protein 697 | 1.825199507 | 4.507781193 | 12.51870781 | 3.64E-12 | 2.47E-09 |
| OSGIN1 | oxidative stress induced growth inhibitor 1 | 1.976862839 | 3.242178076 | 12.4693834 | 3.99E-12 | 2.56E-09 |
| LRP12 | LDL receptor related protein 12 | 1.150991345 | 5.879349667 | 12.33758796 | 4.73E-12 | 2.74E-09 |
| ENPP2 | ectonucleotide pyrophosphatase/phosphodiesterase 2 | 1.132881762 | 4.159837867 | 12.312964 | 4.93E-12 | 2.74E-09 |
| CA12 | carbonic anhydrase 12 | 2.952611083 | 2.055537269 | 12.06586364 | 8.50E-12 | 4.16E-09 |
| GDNF | glial cell derived neurotrophic factor | 2.221923989 | 1.953632019 | 11.88345072 | 1.09E-11 | 4.96E-09 |
| RAPGEF3 | Rap guanine nucleotide exchange factor 3 | 1.358483123 | 5.317230183 | 11.84390589 | 1.10E-11 | 4.96E-09 |
| DNMBP | dynamin binding protein | 1.649697916 | 5.340547806 | 11.79126712 | 1.22E-11 | 5.14E-09 |
| MMP1 | matrix metallopeptidase 1 | 2.614142666 | 4.055963827 | 11.80725757 | 1.28E-11 | 5.14E-09 |
| DOCK4 | dedicator of cytokinesis 4 | 1.442918734 | 5.114520434 | 11.74210078 | 1.31E-11 | 5.14E-09 |
| HMGA2 | high mobility group AT-hook 2 | 1.693818881 | 8.02876654 | 11.73605566 | 1.35E-11 | 5.14E-09 |
| TUBB4A | tubulin beta 4A class IVa | 1.716661008 | 3.085356996 | 11.70972736 | 1.41E-11 | 5.22E-09 |

Table S6 Top 20 downregulated DEGs ranked by adjusted P-value (adj.P.Val) with a logFC of ≤-1 from D9 of the 20 nm PS NPs treatment group. Associated with Fig. 2

| Gene Symbol | Description | logFC | AveExpr | t | P.Value | adj.P.Val |
| --- | --- | --- | --- | --- | --- | --- |
| NTN4 | netrin 4 | -1.889890294 | 5.225618908 | -15.24726237 | 5.77E-14 | 2.35E-10 |
| C1orf116 | chromosome 1 open reading frame 116 | -1.612028614 | 4.917871525 | -13.98298412 | 3.57E-13 | 8.73E-10 |
| PRSS23 | serine protease 23 | -1.723874785 | 8.195470608 | -13.27241203 | 1.07E-12 | 1.88E-09 |
| RBP7 | retinol binding protein 7 | -1.66418135 | 4.895768223 | -13.18407907 | 1.23E-12 | 1.88E-09 |
| PDGFRB | platelet derived growth factor receptor beta | -4.048490407 | 1.053700049 | -13.13275912 | 1.61E-12 | 1.89E-09 |
| POGLUT2 | protein O-glucosyltransferase 2 | -1.755867472 | 7.632600439 | -12.91084107 | 1.91E-12 | 1.95E-09 |
| LINC01111 | long intergenic non-protein coding RNA 1111 | -1.981062476 | 4.82294845 | -12.72611155 | 2.62E-12 | 2.29E-09 |
| FGB | fibrinogen beta chain | -3.745078368 | 6.66080506 | -12.69469838 | 3.18E-12 | 2.31E-09 |
| LEPROT | leptin receptor overlapping transcript | -1.388749124 | 8.267604943 | -12.57785099 | 3.21E-12 | 2.31E-09 |
| RAB11FIP1 | RAB11 family interacting protein 1 | -1.448743991 | 7.508184191 | -12.3474445 | 4.71E-12 | 2.74E-09 |
| SFN | stratifin | -1.499097832 | 5.750216464 | -12.29040081 | 5.20E-12 | 2.76E-09 |
| LMCD1 | LIM and cysteine rich domains 1 | -2.035603782 | 7.197283722 | -12.01954326 | 8.50E-12 | 4.16E-09 |
| DCDC2 | doublecortin domain containing 2 | -1.204645456 | 7.311001135 | -11.80807003 | 1.16E-11 | 5.06E-09 |
| CAV2 | caveolin 2 | -1.073950029 | 7.517799227 | -11.64328235 | 1.53E-11 | 5.24E-09 |
| LRG1 | leucine rich alpha-2-glycoprotein 1 | -2.940359904 | 3.21405824 | -11.63902043 | 1.76E-11 | 5.83E-09 |
| KRT80 | keratin 80 | -1.241892164 | 7.830402556 | -11.40567375 | 2.33E-11 | 7.13E-09 |
| WNT2B | Wnt family member 2B | -2.50384765 | 4.321602236 | -11.43656486 | 2.42E-11 | 7.22E-09 |
| ST6GALNAC5 | ST6 N-acetylgalactosaminide alpha-2,6-sialyltransferase 5 | -1.699983613 | 3.549159648 | -11.38025203 | 2.50E-11 | 7.26E-09 |
| DUSP1 | dual specificity phosphatase 1 | -1.331132104 | 8.359501451 | -11.35653596 | 2.55E-11 | 7.26E-09 |
| TRAM1 | translocation associated membrane protein 1 | -1.216339594 | 8.870800509 | -11.30966056 | 2.76E-11 | 7.67E-09 |

Table S7 Top 3 upregulated DEGs ranked by adjusted P-value (adj.P.Val) with a logFC of ≥1 from D6 of the 100 nm PS NPs treatment group. Associated with Fig. 2

| Gene Symbol | Description | logFC | AveExpr | t | P.Value | adj.P.Val |
| --- | --- | --- | --- | --- | --- | --- |
| ERVH-3 | endogenous retrovirus group H member 3 | 1.190084111 | 4.577821091 | 6.99208763 | 1.80E-07 | 0.000439257 |
| KCNQ3 | potassium voltage-gated channel subfamily Q member 3 | 1.096962388 | 3.050186402 | 5.137950381 | 1.60E-05 | 0.011942982 |
| NYAP2 | neuronal tyrosine-phosphorylated phosphoinositide-3-kinase adaptor 2 | 1.774672497 | 1.062146478 | 4.401971938 | 0.000114464 | 0.045117193 |

Table S8 Top 3 downregulated DEGs ranked by adjusted P-value (adj.P.Val) with a logFC of ≤-1 from D9 of the 100 nm PS NPs treatment group. Associated with Fig. 2

| Gene Symbol | Description | logFC | AveExpr | t | P.Value | adj.P.Val |
| --- | --- | --- | --- | --- | --- | --- |
| IL11 | interleukin 11 | -1.448082119 | 5.610287656 | -7.17739326 | 1.20E-07 | 0.000382735 |
| WNT2B | Wnt family member 2B | -1.21969997 | 4.321602236 | -6.102442038 | 1.49E-06 | 0.003029794 |
| EPPK1 | epiplakin 1 | -1.520538517 | 2.163458516 | -5.219047678 | 1.37E-05 | 0.011942982 |

Table S9 Top 6 hallmark pathways from D6 of the 20 nm PS NPs treatment group and ranked by adjusted P-value (adj.P.Val). Associated with Fig. 4

| pathway | pval | padj | log2err | ES | NES | leadingEdge | time | treatment | regulation | log10padj |
| --- | --- | --- | --- | --- | --- | --- | --- | --- | --- | --- |
| KRAS SIGNALING UP | 0.142191 | 0.544974 | 0.179782 | 0.434985 | 1.319077 | ANGPTL4, TMEM158, CSF2 | D6 | 20 | Up | 0.263625 |
| ESTROGEN RESPONSE LATE | 0.181658 | 0.544974 | 0.135002 | -0.47817 | -1.30649 | EGR3, CCN5, CDH1, FOS | D6 | 20 | Down | 0.263625 |
| APOPTOSIS | 0.467372 | 0.649038 | 0.076084 | -0.3656 | -0.99891 | EGR3 | D6 | 20 | Down | 0.18773 |
| UV RESPONSE UP | 0.527928 | 0.649038 | 0.070961 | -0.37517 | -0.93632 | NR4A1, FOS, ATF3, FOSB | D6 | 20 | Down | 0.18773 |
| TNFA SIGNALING VIA NFKB | 0.614213 | 0.649038 | 0.060509 | -0.26399 | -0.8474 | EGR3, NR4A1 | D6 | 20 | Down | 0.18773 |
| HYPOXIA | 0.649038 | 0.649038 | 0.075118 | 0.271253 | 0.86847 | ANGPTL4 | D6 | 20 | Up | 0.18773 |

Table S10 Top 4 significant hallmark pathways from D9 and overlapping pathways from D6 of the 20 nm PS NPs treatment group. Ranked by adjusted P-value (adj.P.Val). Associated with Fig. 4

| Pathway | pval | padj | log2err | ES | NES | leadingEdge | time | treatment | regulation | log10padj |
| --- | --- | --- | --- | --- | --- | --- | --- | --- | --- | --- |
| KRAS SIGNALING UP* | 0.000106 | 0.003505 | 0.538434 | 0.460477 | 2.409424 | ANGPTL4, TMEM158, EMP1, G0S2, ETV4, PLAT, EPB41L3, CSF2, EREG, DNMBP, PLAUR, ETV5, SOX9 | D9 | 20 | Up | 2.455251 |
| FATTY ACID METABOLISM | 0.00437 | 0.046306 | 0.407018 | 0.560719 | 1.912626 | IL4I1, G0S2, CPT1A, ACSL1, ACSL5 | D9 | 20 | Up | 1.334361 |
| G2M CHECKPOINT | 0.004597 | 0.046306 | 0.407018 | 0.689133 | 1.913909 | CCND1, HMGA1, ODC1, ATF5, TLE3, SLC12A2 | D9 | 20 | Up | 1.334361 |
| IL2 STAT5 SIGNALING | 0.005613 | 0.046306 | 0.407018 | 0.402031 | 1.921216 | EMP1, NT5E, ETV4, CSF2, SPRY4, TNFRSF8, PHLDA1, PLIN2, ODC1, CDCP1, FAH | D9 | 20 | Up | 1.334361 |
| ESTROGEN RESPONSE LATE* | 0.057096 | 0.356218 | 0.24134 | -0.35135 | -1.55129 | CCN5, EGR3, CDH1, FOS, PRSS23, GJB3, NPY1R, SEMA3B, ALDH3B1, OPN3, SFN | D9 | 20 | Down | 0.448285 |
| HYPOXIA* | 0.153652 | 0.489905 | 0.179782 | 0.276747 | 1.356299 | ANGPTL4, HMOX1, CA12 | D9 | 20 | Up | 0.309888 |
| UV RESPONSE UP* | 0.15894 | 0.489905 | 0.140406 | -0.34859 | -1.32689 | FOS, ATF3, JUNB, ABCB1, CDKN1C, NR4A1, FOSB, KCNH2, RASGRP1 | D9 | 20 | Down | 0.309888 |
| APOPTOSIS* | 0.176948 | 0.489905 | 0.130777 | -0.3259 | -1.29774 | PDGFRB, EGR3, ERBB3, F2, ATF3 | D9 | 20 | Down | 0.309888 |
| TNFA SIGNALING VIA NFKB* | 0.574344 | 0.74249 | 0.09226 | 0.167767 | 0.913155 | G0S2, CCND1, CSF2, PTX3, PLAUR, PHLDA1, AREG, LAMB3, IL18, MAP2K3, TNC, IL7R, TNFSF9, CSF1, ATP2B1 | D9 | 20 | Up | 0.129309 |
| * Pathway present at D6 | | | | | | | | | | |

#### Supplementary Figures

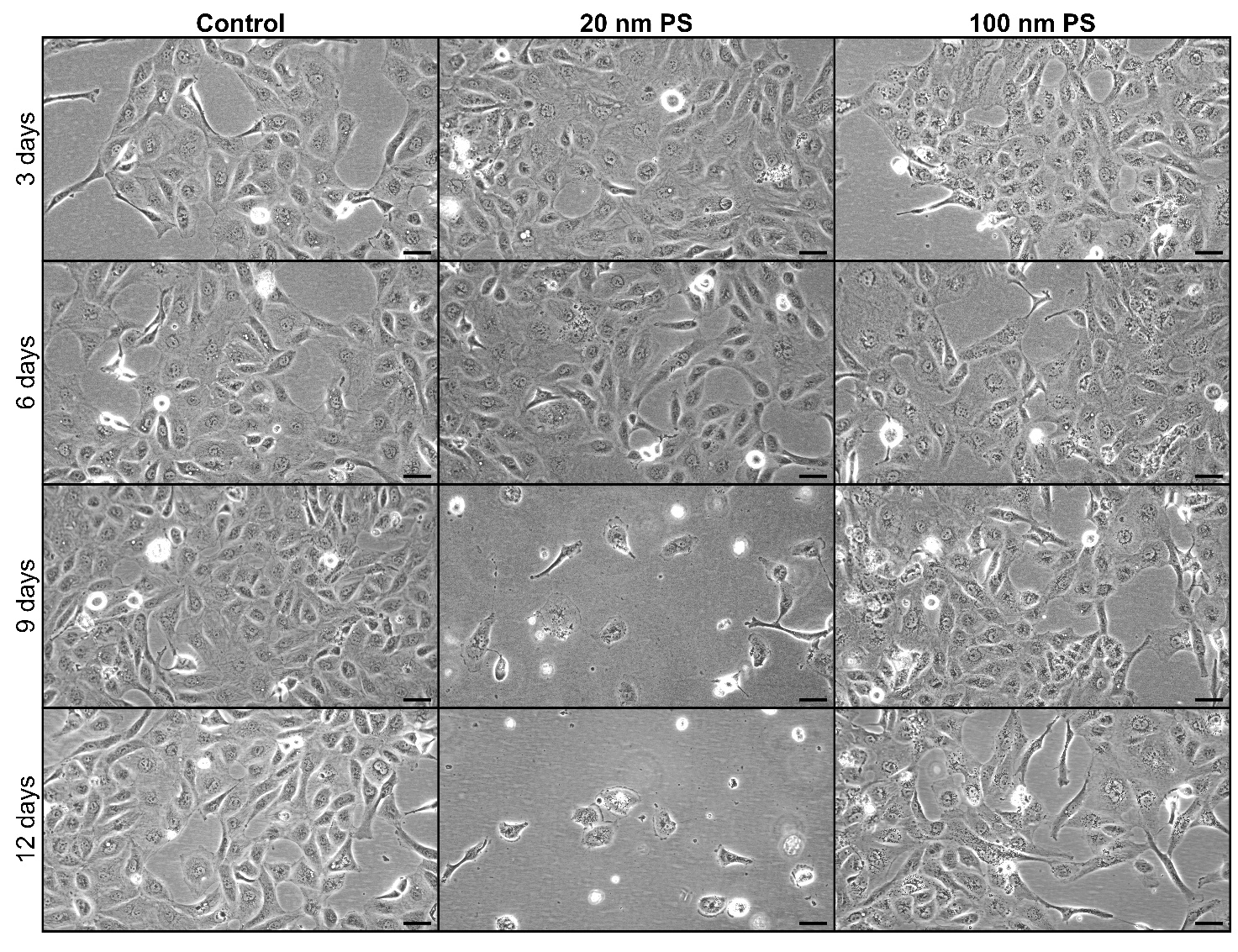

Fig. S1 Micrographs of HK-2 cells following repeated exposure to 100 µg/mL 20 nm and 100 nm PS NPs. All cells were treated with either PS20-A, PS100-A, PS15-B or PS100-B NPs at increasing concentrations from 0.2 µg/mL to 200 µg/mL, top to bottom. 20x magnification. Scale bar = 50 µm. N=3. Data is representative of individual replicates tested

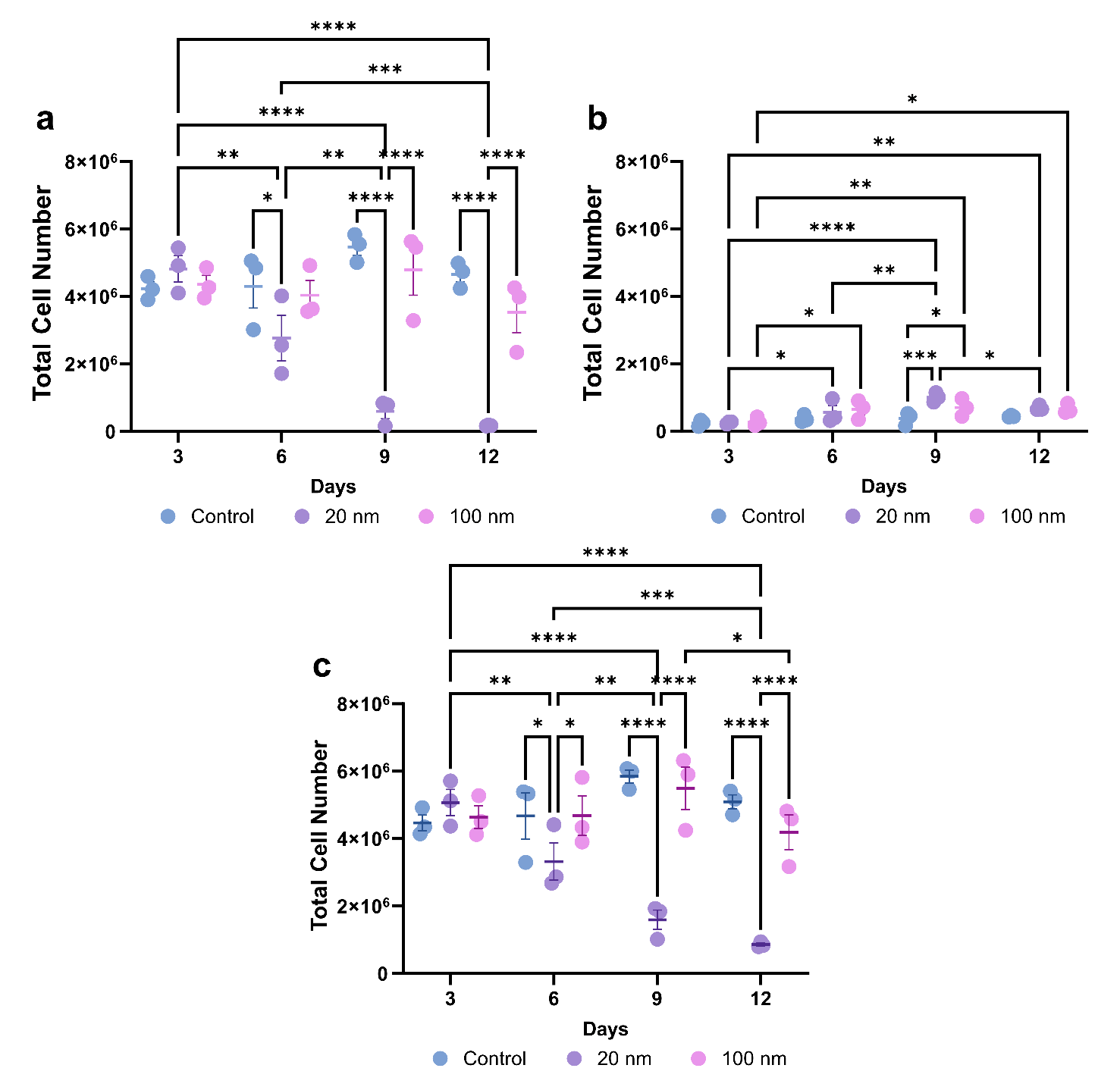

Fig. S2 a) Total live cell number of HK-2 cells following chronic, repeated exposure to PS NPs. Cells were grown for a total 72-hours between splits. b) Total dead cell number of HK-2 cells after repeated exposure to 20 nm or 100 nm PSNPs over 12 days. c) Total cell number of HK-2 cells after repeated exposure to 20 nm or 100 nm PSNPs over 12 days. P-value = * ≤0.05, ** ≤0.01, *** ≤0.001, **** ≤0.0001.

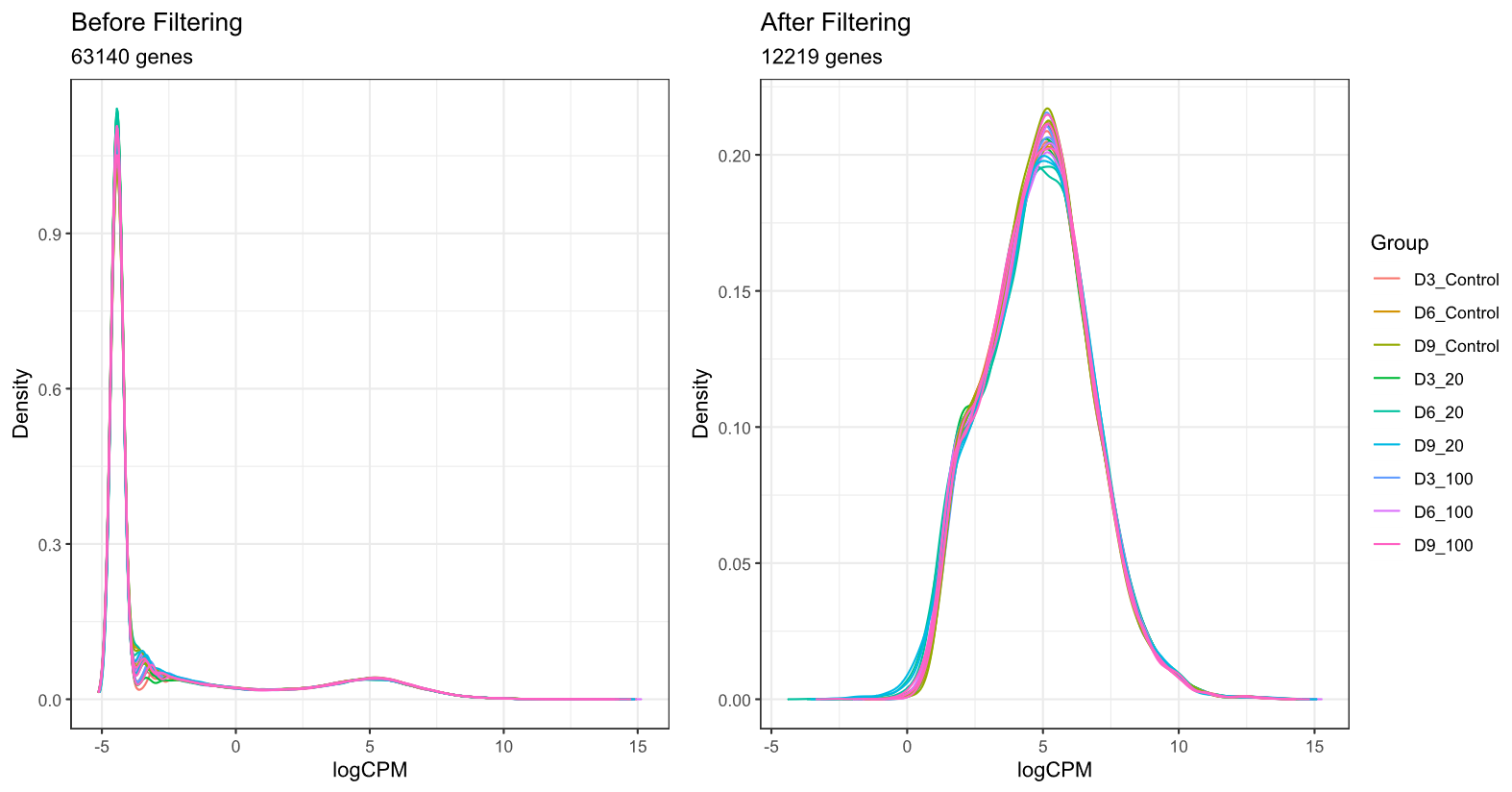

Fig. S3 Density distribution of sample genes pre- and post-filtering.

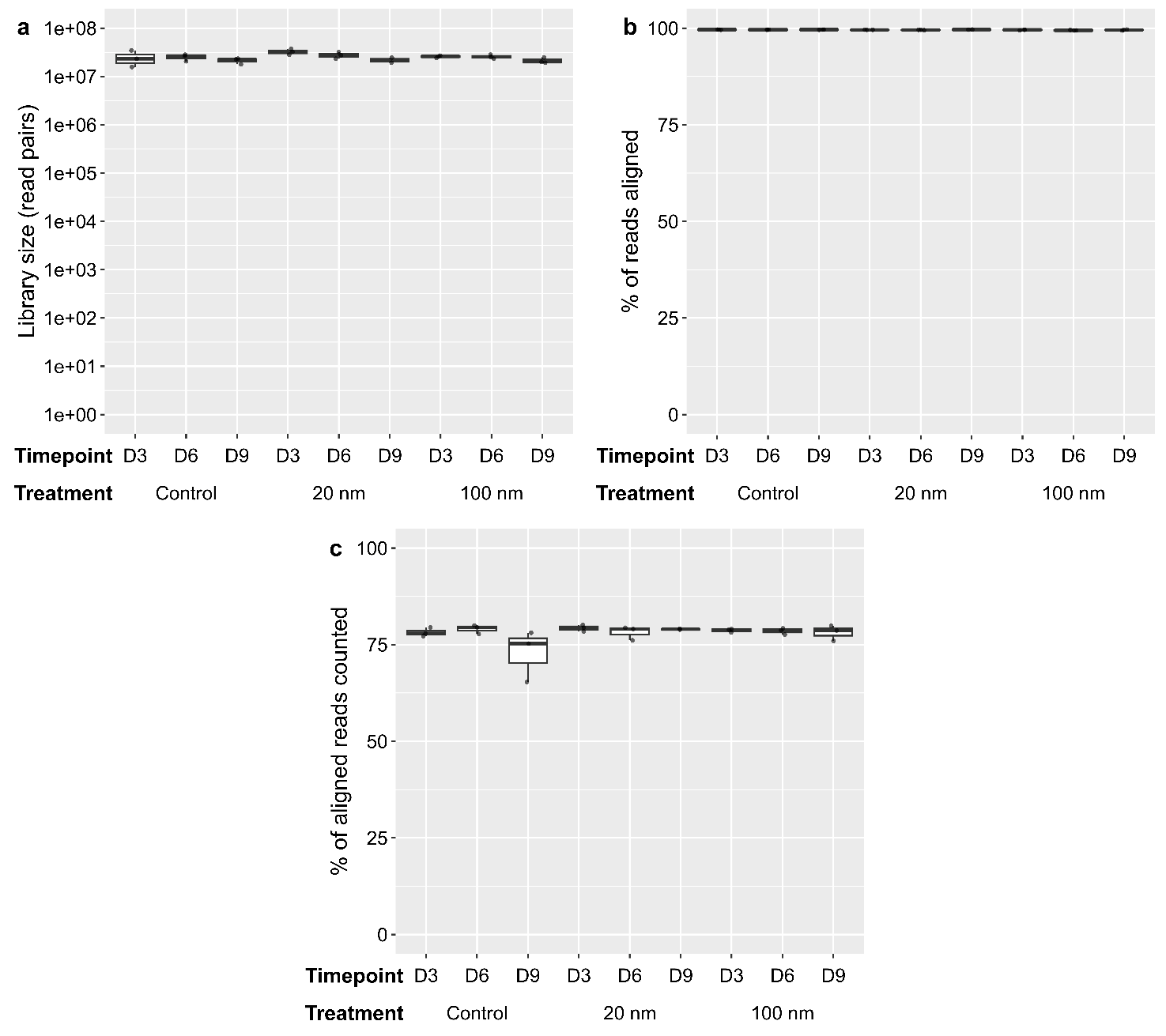

Fig. S4 Library QC and MDS alignment of samples. a) Library size distribution between samples. b) Percentage of aligned reads between samples. c) Percentage of counted aligned reads.

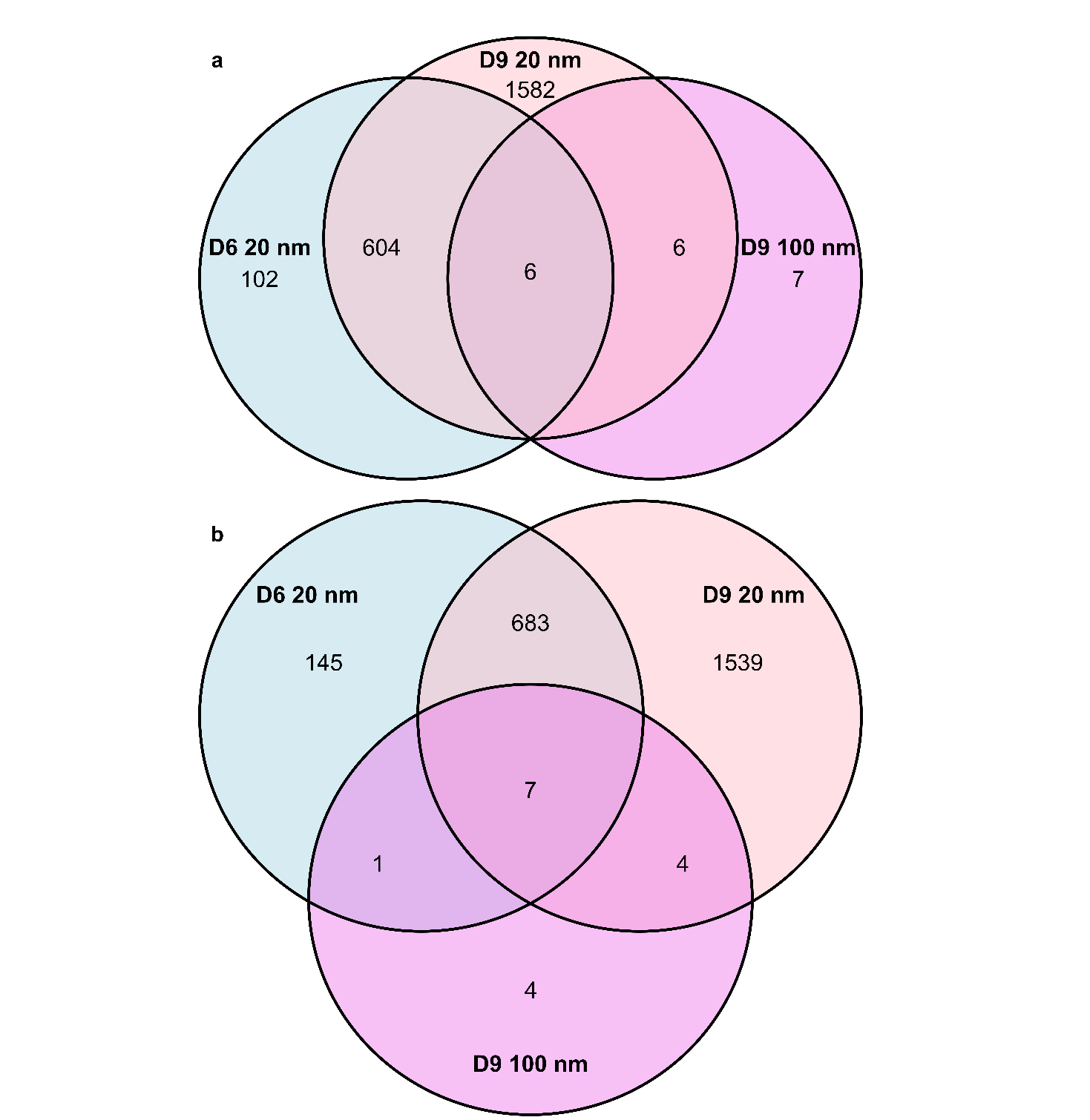

Fig. S5 Venn diagrams of the overlap in a) upregulated and b) downregulated DEGs between the D6 and D9 20 nm PS NPs and D9 100 nm PS NPs treatment groups.

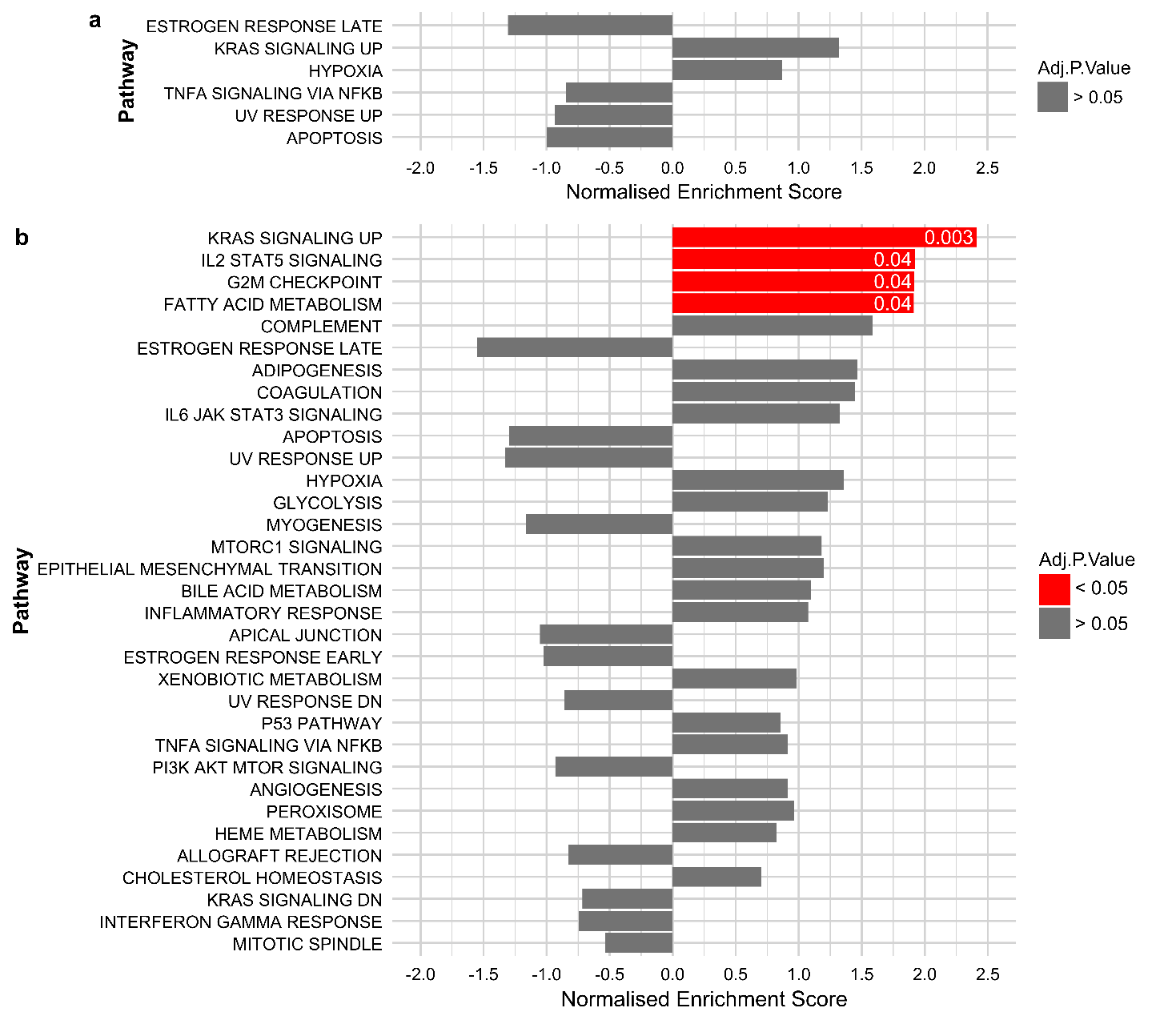

Fig. S6 Full hallmark pathway analysis outputs. a) D6 20 nm. b) D9 20 nm.
